# Multi-trait GWAS for diverse ancestries: Mapping the knowledge gap

**DOI:** 10.1101/2023.06.23.546248

**Authors:** Lucie Troubat, Deniz Fettahoglu, Léo Henches, Hugues Aschard, Hanna Julienne

## Abstract

**Background:** Approximately 95% of samples analyzed in univariate genome-wide association studies (GWAS) are of European ancestry. This bias toward European ancestry populations in association screening also exists for other analyses and methods that are often developed and tested on European ancestry only. However, existing data in non-European populations, which are often of modest sample size, could benefit from innovative approaches as recently illustrated in the context of polygenic risk scores.

**Methods:** Here, we extend and assess the potential limitations and gain of our multi-trait GWAS pipeline, **JASS** (**J**oint **A**nalysis of **S**ummary **S**tatistics), for the analysis of non-European ancestries. To this end, we conducted the joint GWAS of 19 hematological traits and glycemic traits across five ancestries (European (EUR), admixed American (AMR), African (AFR), East Asian (EAS), South-East Asian (SAS)).

**Results:** We detected 367 new genome-wide significant associations in non-European populations (15 in Admixed American (AMR), 72 in African (AFR) and 280 in East Asian (EAS)). New associations detected represent 5%, 17% and 13% of associations in the AFR, AMR and EAS populations, respectively. Overall, multi-trait testing increases the replication of European associated loci in non-European ancestry by 15%. Pleiotropic effects were highly similar at significant loci across ancestries (e.g. the mean correlation between multi-trait genetic effects of EUR and EAS ancestries was 0.88). For hematological traits, strong discrepancies in multitrait genetic effects are tied to known evolutionary divergences: the ARKC1 loci, which is adaptive to overcome the *p.vivax* induced malaria.

**Conclusions:** Multi-trait GWAS can be a valuable tool to narrow the genetic knowledge gap between European and non-European populations.

## Background

After 15 years of intensive GWAS analyses, investigators are reaping tangible translational benefits such as drug repurposing opportunity, and actionable polygenic risk scores [1]. Yet, ∼95% of samples analyzed in GWAS are from European ancestry [2]. Genetic insights gained on European ancestry are only partially transferrable to other ancestries: polygenic risk scores have lessened accuracy [3–5], and genetic testing yields more often ambiguous results in non-European ancestries [6]. Altogether, this gap in data and analyses is a major bias in the existing research that can result in increased health disparities [7,8]. The genetic community is increasingly aware of this issue and strongly recommend to include more individuals from non-European ancestries in GWAS studies [2,7,9–12]. Increasing the sample size coverage of non-European ancestries has been identified as one of the most important objectives for the genetic community in the coming years [1].

Increasing the sampling of non-European ancestries is paramount, but computational method development is also an area of improvement that is less often discussed [10,11]. Many statistical genetics methods are developed and tested primarily on the European population. To ascertain this bias, we surveyed 25 recent methods in statistical genetics (Table S1). Amongst these methods 20 conducted an application on human real data, all included European data, and 13 focused on European data exclusively. When developed on European data exclusively, using these computational methods on European ancestry becomes all the more easy: input files (e.g. specific formats of reference panels) are often provided, performances and potential pitfalls are known, and protocols (e.g. parameters) are specific. In comparison, applying these methods to diverse ancestries can require substantial additional work and can be perceived as riskier since potential performance discrepancies are undocumented.

This *European by default* mode creates a snowballing effect where secondary analysis increases further the genetic knowledge gap between European and non-European ancestries. Furthermore, the analysis of non-European do not only address a public health gap, but also offer opportunities and several recent publications illustrate the benefit of methodological development for the analysis of diverse populations in various applications such as: polygenic risk scores [13], association testing [14], and TWAS [15].

In this context, we tested if the properties of our previously developed JASS (Joint Analysis of Summary Statistics) pipeline – its ability to detect biologically relevant associations missed by univariate GWAS [16,17] – would generalize to non-European ancestries. To this end, we conducted the joint GWAS of 19 hematological traits and glycemic traits retrieved from two large trans ancestry studies [18,19]. We applied our pipeline to five ancestries (European (EUR), Admixed American (AMR), African (AFR), East Asian (EAS), South-East Asian (SAS)), and investigated the performance and robustness of each step of the pipeline across ancestries. For the given input data, the pipeline generalized properly to three out of the four non-European ancestries: AMR, AFR, and EAS. We detected 367 new genome-wide significant associations in non-European populations (15 in AMR, 72 in AFR and 280 in EAS). We then report to what extent multi-trait testing increases the replication of European associated loci in non-European ancestry. Finally, we investigate the conservation of pleiotropy across ancestries, and tie the observed differences to known evolutionary divergences. Altogether these analyses suggest that multi-trait GWAS methods can be valuable to take advantage of modest size cohorts and to narrow the genetic knowledge gap between European and non-European populations.

## Methods

### Survey of ancestries used in recently published methods in statistical genetics

To ascertain a potential bias in ancestries used to develop and test recent statistical genetic tools, we surveyed 25 publications. We looked up ancestries studied in: i) methodology focused publications of the *American Journal of Human Genetics* published after January 2023, ii) *Bioinformatics* publications in the “Genetics and Population Analysis” section of the March, April and May issues, and iii) the first page of results of the Google Scholar query “gwas summary statistics methods” ordered by relevance and published after 2021 (query was performed by Dr. Julienne on May 17^th^ 2023). This survey does not pretend to comprehensively ascertain the ancestry bias in method development in statistical genetics. Its aim is rather to provide a snapshot of common practices of the field at the time of this study.

### Summary Statistics

GWAS of hematological traits originate from the Chen et al paper [19] and were downloaded from the GWAS Catalog (https://www.ebi.ac.uk/gwas/publications/32888493#study_panel). We analyzed 15 quantitative hematological traits in 740,292 participants from four ancestries: 563 946 Europeans, 151,807 East Asians, 9,368 Admixed Americans and 15 171 Africans (Table S2). GWAS of glycemic traits come from the [18] study downloadable from GWAS Catalog (https://www.ebi.ac.uk/gwas/publications/34059833). We analyzed four glycemic traits in 281,416 individuals without diabetes from European, East Asian, Admixed American and African American or Afro-Caribbean (African population) populations (Table S2). Links towards harmonized, imputed, and curated summary statistics using the JASS pipeline are provided in the **“Availability of data and material”** section.

### Data preprocessing

We leveraged our previously published method, JASS[16,17], to harmonize data of each ancestry for multi-trait GWAS. Briefly, the JASS pipeline is implemented in nextflow and proceeds through four preprocessing steps: i) mapping and alignment on the reference panel, ii) harmonization of the sample size, iii) computation of the genetic covariance matrices using the LDScore regression and, iv) imputation. In addition to summary statistics, the pipeline requires the following input for each ancestry: a reference panel, a file indicating LD independent regions, LD matrices for imputation (using Robust and Accurate imputation from Summary Statistics, RAISS) and input files for the LDSC regression[20].

Input files for the JASS pipeline were prepared for the five ancestries. A reference panel for common SNPs (MAF > 1%) was built from 1000 Genomes consortium[21] phase 3 data (hg38 build). When absent in 1000G, an rsid identifier was retrieved from all germline variations listed in Ensembl (http://ftp.ensembl.org/pub/release-105/variation/gvf/homo_sapiens). LD score files were computed using the ldsc.py script available at https://github.com/bulik/ldsc with the following inline options: --l2 --ld-wind-kb 1000. LD matrices for imputation were generated using the generate_genome_matrices function from the RAISS python package (see https://statistical-genetics.pages.pasteur.fr/raiss/#precomputation-of-ld-correlation).

RAISS Imputation parameters were chosen to ensure high accuracy for all traits and ancestries as described in the documentation (https://statistical-genetics.pages.pasteur.fr/raiss/#optimizing-raiss-parameters-for-your-data). We systematically assessed RAISS accuracy for all traits and ancestries on chr22. We set RAISS key parameters to --R2-threshold 0.6, --minimum-ld 5 and --eigen-threshold 0.05 and report corresponding accuracies in Table S3. The correlations between the imputed and initial values range from 0.93 to 0.99.

Links towards the JASS pipeline and associated computational tools are provided in the “Availability of data and material” **section.**

### LD independent regions

For each ancestry, we computed LD independent regions using the R package bigsnpr [22] designed for massive analysis of SNP arrays. The function snp_ldsplit splits a correlation matrix in quasi-independent blocks using dynamic programming [23]. We optimized settings to reach a tradeoff between the sum of squared correlations outside the blocks (cost) and the number of regions found in a chromosome (n_block). On the chromosome 22 of each ancestry, we have varied the minimum number of variants in each block (min_size) between 250 and 10 000, the maximum number of variants in each block (max_size) between 1000 and 50 000, and the threshold under which squared correlations are ignored (thr_r2) between 0.005 and 0.4. By analyzing the results, we selected thr_r2 = 0.05, min_size = 4000, max_size = 5000, max_K = 400 as it provides a balance between the cost and the number of regions for all chromosomes of all ancestry (Figure S1).

### The omnibus multi-trait test and contrast with univariate GWAS

To perform the *omnibus* multi-trait tests, JASS relies on a theoretical framework and a software architecture described in [16,17]. Briefly, assuming an infinitesimal model and under the null hypothesis of no genetic effects, the statistic of the omnibus test (equation 1) follows *a X^2^(k)* distribution where denotes the number of traits considered in the study. The omnibus test statistic is defined as:

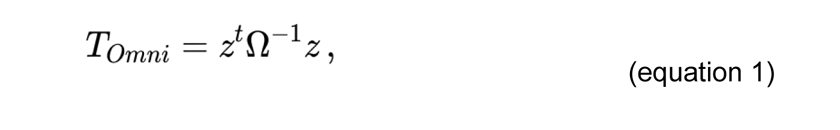

where is the vector of the z-score across traits and Ω is the residual z-score covariance under the null hypothesis. As derived in [20], for a pair of traits i and j the expected covariance under the null hypothesis is equal to 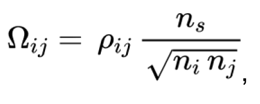, where is the total covariance between traits and *pij*, is the number of samples shared between studies and *j*, and and are the sample sizes of studies and *j*.

### Validation of the estimation of *Ω* by the LDscore regression in non-European ancestries

To ensure that the *Ω*parameter of the omnibus test is correctly estimated in non-European ancestries, we compared the direct estimate of *pij* computed in UK Biobank and an indirect estimate derived from the LDscore regression applied to GWAS summary statistics. We used the ancestry segmentation presented in [4] and worked with the Indian (equivalent to SAS), Nigerian (equivalent to AFR), and Caribbean (equivalent to AFR with admixture with EUR) individuals. We restricted our analysis to complete cases (Sample sizes of 5565, 3427, and 1959 for Indian, Nigerian, and Caribbean individuals respectively) for hematological traits so the term *ns/√n_1_n_2_* equals 1. For each population, we derived a direct estimate of by computing the Pearson correlation between hematological traits in UK BioBank. We then derived LDscore estimates by: i) computing univariate GWAS on each trait using plink, ii) applying the JASS pipeline on the resulting summary statistics. To compute the LDscore for each variant, we used the SAS 1000G reference panel for the Indian population and the AFR 1000G reference panel for the Caribbean and Nigerian populations (**Availability of data and material**).

### Characterization of new genetic associations

JASS identifies significant genetic variants using a two step procedure. First, large LD independent regions (see paragraph above for region computation) are scanned for new associations. For each LD independent region, the minimum association p-value for the univariate test across traits and for the joint test are retrieved. In regions containing significant association for JASS, we searched for potential additional lead SNPs using the PLINK clump function [24] with the following parameters: a significance threshold equal or greater than 5.10^-8^ (--clump-p1), a significance threshold for other SNPs in the clump equal or greater than 5.10^-4^ (--clump-p2) and a linkage disequilibrium of *r*^2^=0.2 (--clump-r2). We filtered out isolated lead SNPs (clumps containing only the lead SNPs) to remove poor quality associations. A lead SNPs is considered as a new association if the p-value of the omnibus test is significant while the minimum univariate test p-value across traits is not. Note that we choose to apply no multi testing correction to the minimum univariate p-value as we wish to contrast JASS results with all associations reported previously in the literature.

### Trans ancestry meta analysis

We performed a trans ancestry multi-trait GWAS by adapting the omnibus test to the trans ancestry setting. Under the null hypothesis, the omnibus statistic follows a *X^2^(k)* distribution with *k* degree of freedom where *k* denotes the number of traits considered. As a cohort of a given ancestry is independent from cohorts of other ancestries, the sum of the omnibus statistics across all ancestries, *A* = {*AFR; EUR; EAS; AMR*}, follows a chi-square with *k* × (*#ancestries*) degree of freedom.

For a given SNPs:

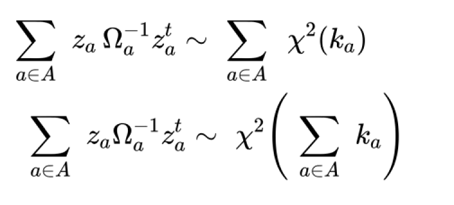

where *z* is the vector of z-scores across traits, *A* is the ancestry considered and *Ω^A^* is the residual covariance between Z-score under the null for the ancestry *A*.

To contrast the multi-trait trans ancestry analysis against univariate meta-analysis we performed a meta-analysis on Z-score (assuming fixed genetic effects) and retrieved the minimum p-value across traits.

For a given trait, we computed the trans ancestry meta analysis Z-score as:

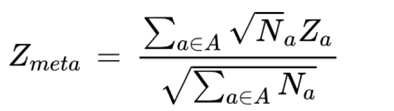

As LD independent regions cannot be defined across ancestries, we define regions as 1Mbp blocks and select one lead SNPs per block as the one with a minimum p-value.

### Simulating a smaller sample size for European data

To simulate a diminished sample size for the European data, we scaled down Z-scores of lead SNPs with the following formula:

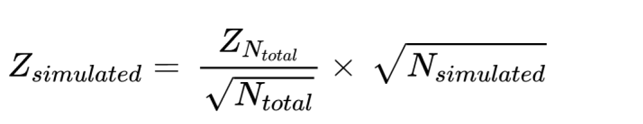

Once scaled down, new p-values for the univariate test and the omnibus test were computed. We varied the simulated sample size from a fraction of 0 to 100% of the initial sample by increments of 1%.

### Functional annotation

We mapped lead SNPs to genes through positional mapping and eQTLs (expression Quantitative Trait Loci) using Functional Mapping and Annotation of Genome-Wide Association Studies (FUMA GWAS) [25]. For position based mapping, SNPs were mapped to their nearest gene if the distance was under 10kb. For eQTL mapping, we selected blood and immune system tissues (eQTL catalog (BLUEPRINT monocyte, BLUEPRINT neutrophil, BLUEPRINT T-cell, CEDAR platelet, CEDAR T-cell CD4, CEDAR T-cell CD8, GENCORD T-cell, Kasela 2017 T-cell CD4, Kasela 2017 T-cell CD8, Lepik 2017 blood, Naranbhai 2015 neutrophil CD16), van der Wijst et al. scRNA eQTLs (B cells, Dendritic cells, Natural Killer cells, Monocytes, Classical Monocyte, Non-classical Monocytes, CD4 T cells, CD8 T cells, PBMC), DICE (Naive B cells, Naive CD4 T cells, Naive (activated) CD4 T cells, Naive CD8 T cells, Naive (activated) CD8 T cells, Classical Monocytes, Non-classical Monocytes, Natural Killer cells, TFH CD4 T cells, TH1 CD4 T cells, TH17 CD4 T cells, TH1-17 CD4 T cells, TH2 CD4 T cells, Memory TREG CD4 T cells, Naive TREG CD4 T cells), Blood eQTLs (Westra et al. (2013) BIOS QTL Browser), GTEx v8 Blood Vessel (GTEx Cells EBV-transformed lymphocytes, GTEx Whole Blood)).

To compute the genomic distance between novel associations mapped to the same genes in the EUR and EAS ancestries, we retrieved genes linked by significant eQTLs to SNPs discovered by JASS exclusively in each ancestry. Then, using the intersection of EAS and EUR genes, we computed the genomic distance between the linked SNP in EAS and the linked SNP in EUR for each gene. If a gene was linked to multiple SNPs, the pair of SNPs with the smallest distance was retained.

For the investigation of newly detected genes in African ancestry, we focused on genes tied to SNPs discovered by JASS exclusively and belonging to at least one GeneOntology (GO) set. Then, we queried each of this gene function on The Human Gene Database GeneCards (www.genecards.org, [26]).

## Results

### Overview of the study

Our analysis focused on 15 hematological and four glycemic quantitative traits in five superpopulations (**Figure 1**). We retrieve summary statistics from two large trans ancestry meta-analyses each focusing either on hematological traits or on glycemic traits. Using a single source for summary statistics by group of traits allows to minimize potential discrepancies in data preprocessing across ancestries. It also allows for an increased sample overlap between traits, which can be beneficial for the statistical power of the multi-trait test[16,27,28]. Sample size varies widely between populations: GWAS on hematological phenotypes studied 563,946 individuals from EUR ancestry, 151,807 for the EAS ancestry, 9,368 for AMR ancestry, 15,171 for AFR ancestry, and 8189 for SAS ancestry (**Table S2**). While the sample size across hematological traits is relatively homogeneous, the sample size in glycemic traits presents large discrepancies. For instance, in the AFR population, ten times more individuals were analyzed for fasting glucose phenotype than for the two-hour glucose tolerance test. We applied a previously developed pipeline [17] to: i) curate and harmonize GWAS summary statistics, ii) impute missing statistics within each study using the RAISS software (Robust and Accurate Imputation from Summary Statistics,[29]), and iii) compute multi-trait GWAS. Entry files for the Nextflow pipeline were generated and made publicly available to facilitate the analysis of summary statistics from all ancestries (**Supplementary Note 1** and **Availability of data and material**).

**Figure 1.**
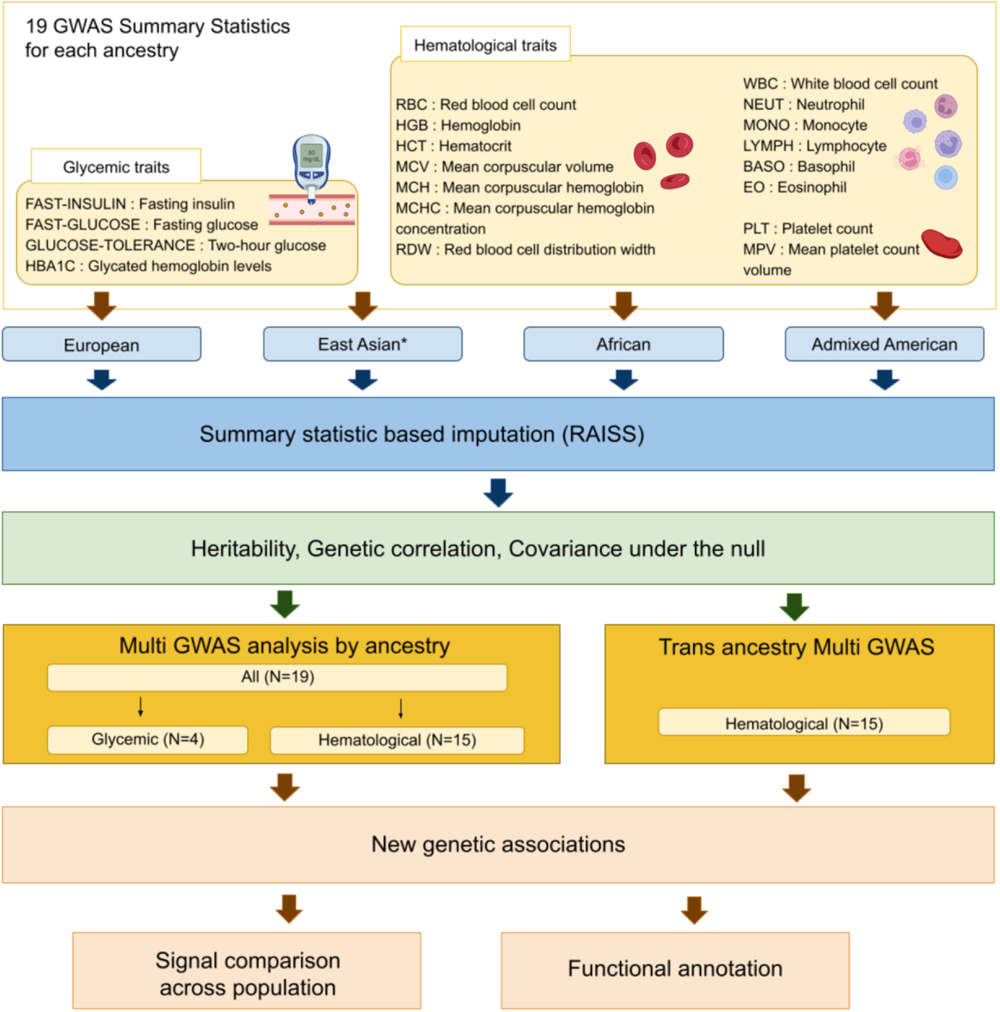
Overview of the study. After harmonization and imputation of GWAS summary statistics, multi-traits GWAS were performed for the four ancestries on hematological and glycemic phenotypes separately. We computed the heritability, genetic correlation and association statistics covariance under the null for each ancestry. An additional trans-ancestry multi-trait GWAS was conducted for hematological traits. Significant hematological associations were further analyzed through functional annotation and by comparing their position and features across ancestries. *For the EAS ancestry, two traits are missing: mean platelet count volume and red blood cell count distribution width.

The median increase in the number of variants across traits after imputation is 6%, 13%, 22%, 21% and 27% for AFR, AMR, EUR, SAS and EAS ancestries, respectively (**Table S2** and **Figure S2**). The imputation method relies on the linkage disequilibrium (LD) structure; the number of variant increases seems to reflect the differences in LD block length across ancestry[8,21]. Imputation performances and the impact of imputation on the number of significant loci with the univariate test are congruent with our previous observations [29] (**Table S3**). All traits available by ancestry were analyzed jointly using an omnibus test implemented in JASS[16,17] (applied to association z-scores), grouping them by hematological traits and glycemic traits. We used the LDscore regression [20] to estimate the covariance of z-scores under the null hypothesis *Ω*(**Table S4**) – a required input for the omnibus test (**Methods**) – along with heritabilities, and genetic correlation across all traits (**Supplementary note 2, Figure S3** and **Table S5**). The covariance of z-scores under the null (*Ω*) hypothesis is a critical parameter for the validity of the omnibus test. We previously validated its estimation by the LDscore regression in the European population[16]. We reproduce a similar validation in non-European ancestries present in UK biobank (namely the Indian, Nigerian and Caribbean populations[4]). Briefly, when all samples are shared between phenotypes, *Ω*is equal to the Pearson correlation matrix of phenotypes. If one has access to individual level data, the Pearson correlation matrix can be compared to its estimation by the LDscore regression (Methods). Overall, LDscore estimates were accurate (mean absolute error = 0.03) even in the Caribbean population, which is admixed between the European and African ancestries[4] (**Figure S4**).

Overall, all analyses passed standard quality controls except for the SAS ancestry. More precisely, we observed an inflation of the joint test statistics on hematological traits after imputation. This limited robustness seems to arise from the higher collinearity of the residual covariance matrix for this ancestry. We discuss this issue in depth in the supplementary material (**Supplementary note 3** and **Figure S5**). Because we cannot rule out additional sources of biases, we decided to exclude both the multi-trait GWAS on hematological traits and on glycemic traits for the SAS ancestry from subsequent analyses.

Using the intersection of available SNPs across ancestries, we first performed a trans-ancestry multi-trait GWAS by summing the *omnibus* test statistics across ancestries and by contrasting its results with a univariate trans-ancestry meta analysis (see **Methods**). For glycemic traits, the set of SNPs available after performing the intersection was too small (#Nsnps = 8727). Hence, the trans-ancestry GWAS was performed only on hematological traits. Focusing on hematological traits, we compared association positioning and features across ancestries, and systematically studied to what extent loci are shared across ancestries. We assessed the biological relevance of significant associations by annotating them with Functional Mapping and Annotation of Genome-Wide Association Studies (FUMA GWAS,[25]) and The Human Gene Database GeneCards (www.genecards.org,[26]).

### New genetic associations

We report 2,648 new independent associations (**Table 1, Tables S6-S10, Figures S6-S10**). Genomic inflation factors are congruent with an adequate control of the type 1 error ranging from 0.99 for the AMR ancestry to 1.23 for the European ancestry (**Figures S11-S13**). New associations are LD-independent associations significant exclusively for the joint test (**Methods**). The univariate p-value is defined as the minimum p-value of univariate tests across traits. Although this choice leads to an inflation of the univariate p-value, it allows for the inclusion of all associations previously reported in the literature and provides a stringent definition of new associations detected by JASS. For the TRANS ancestry analysis, we derived a TRANS ancestry multi-trait association statistic by calculating the sum of the omnibus test statistics across ancestries. Since samples are independent from one ancestry to another, the obtained statistic follows a *aX^2^_k_* with degrees of freedom equal to the number of ancestries times the number of traits (**Methods**). To contrast this TRANS ancestry multi-trait analysis with a univariate approach, we computed a standard meta-analysis for each trait, assuming fixed genetic effects across ancestries (**Methods**).

**Table 1.** Independent genetic associations detected with the JASS omnibus test using the standard p-value threshold of 5.10^-8^. Number of associations found for 15 blood and four glycemic traits in four different populations with the joint test and the univariate tests. Associations that are significant only with the joint test (new associations) are distinguished from associations that are significant with univariate tests.

|  | #Associations found by | European | East Asian | African | Admixed American | Trans | Total by Test |
| --- | --- | --- | --- | --- | --- | --- | --- |
| Hematological traits | joint test only | 2,173 | 271 | 72 | 14 | 50 | 2,580 |
|  | univariate test and joint test | 14,153 | 981 | 875 | 43 | 1,457 | 17,509 |
|  | univariate test only | 4,800 | 389 | 420 | 53 | 293 | 5,955 |
|  | Total by ancestry | 21,126 | 1,641 | 1367 | 110 | 1,800 | 26,044 |
| Glycemic traits | joint test only | 58 | 9 | 0 | 1 |  | 68 |
|  | univariate test and joint test | 371 | 32 | 0 | 8 |  | 411 |
|  | univariate test only | 89 | 6 | 1 | 0 |  | 96 |
|  | Total by ancestry | 518 | 47 | 1 | 9 |  | 575 |

New associations represent 10%, 17%, 5%, 13%, and 2.8% of associations detected by the joint test in the EUR, EAS, AFR, AMR and TRANS GWAS, respectively. The vast majority of associations are associated with hematological traits (97%), following the same trend as univariate association (97.5% of univariate associations are also associated with hematological traits). JASS identified new genetic associations in modest sample size cohorts (i.e 9,368 individuals for AMR) on hematological traits emphasizing the interest in using non-European data when conducting secondary analyses even when they are seemingly underpowered [10]. Concerning glycemic traits, the very modest number of new signals for AFR and AMR might be explained by the low coverage of glycemic summary statistics for these ancestries (**Table S2**). Indeed, for these two ancestries, the approximate number of SNPs available for glycemic traits is 2 million whereas more than 8 million SNPs were available for hematological traits. Here, we chose to report all associations below the standard p-value threshold of 5e10^-8^ for completeness. However, a stricter choice would have been to apply a Bonferroni correction taking into account the number of multi-trait GWAS performed in the current report (11). When applying a Bonferroni correction, the number of new associations is 41, 6, 169, 1479 and 36 in the AFR, AMR, EAS, EUR and TRANS ancestry analyses respectively.

Focusing on hematological trait associations, 38% of new associations detected in the African population arise from absent or rare variants (minor allele frequency [MAF] < 1%) in other studied populations (**Table S11**). In contrast, 4% and none of the new associations detected in the East Asian or European populations were rare or absent in all other populations. The African population’s unique allele diversity appears to be a key factor in explaining the number of associations discovered in this population.

### Shared loci across ancestries for hematological traits

We compared association loci across populations to assess the extent of their potential overlap across ancestries. To this end, the genome was segmented into non-overlapping regions of 1 Mbp that were considered detected in one ancestry if the region contained at least a significant association (joint or univariate). We represented (Figure 2A) the overlap of loci across populations with an UpSetR plot – an improved Venn diagram for the visualization of numerous set intersections. When including several ancestries, we considered the intersection as newly discovered if at least one of the genetic associations was found by the joint test only (Figure 2A). The joint test increased the discovery of non-European ancestry specific loci by over 33% (from 10 regions to 15 regions). Of these specific loci, 12 regions on 15 were detected in the AFR ancestry and 4 of those associations were discovered exclusively by the joint test. This result is in line with previous reports that African cohorts tend to yield more associations by sample than European cohorts [8]. The large number of specific loci detected in European ancestry samples likely reflects the larger sample size of the European cohorts. Specific loci in the European population tended to have a weaker signal (i.e., higher p-value) than those shared by at least one other ancestry (rank test p-value = 0.008, **Figure S14**). These weaker signals might be detected in other ancestries when reaching an equivalent sample size. To test this hypothesis, we artificially weakened the signal of lead SNPs detected in the European population to simulate a range of diminished sample sizes (**Methods**) and computed the fraction of 1 Mbp loci that remained specific to European ancestry (**Figure S15**). When the sample sizes were similar in the pair of ancestries, the fraction of loci specific to European ancestry ranged from 0 (when compared to AMR) to ∼15% (when compared to EAS). This observation is in line with most genetic associations sharing the same region within different populations as observed in previous studies [30–32].

**Figure 2.**
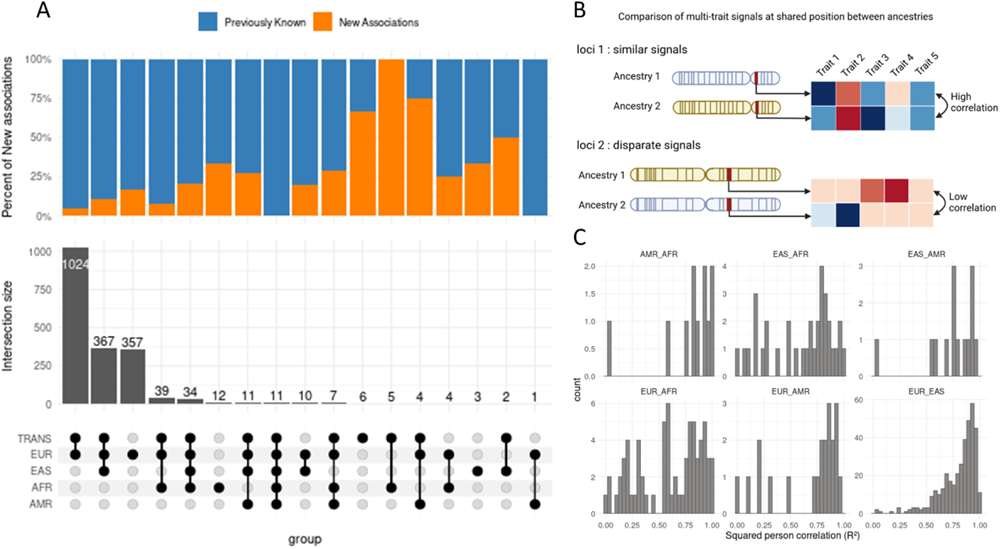
Genetic signal similarity across 4 superpopulations. A) Overlapping loci across four superpopulations. Shows shared and specific loci in the studied populations for hematological traits, B) Illustration of the comparison of multi-trait signals across ancestries. For shared loci a correlation between the vectors of the genetic signal is computed. The squared Pearson correlation gives an indication of how much the multi-trait signal is conserved. C) Histogram of multi-trait genetic signal similarity (R² Pearson correlation) for pairs of lead SNPs located in a shared region. Each panel corresponds to one ancestry pair.

Performing the multi-trait GWAS increased the proportion of significant associations detected in the trans-ancestry and European GWAS that were also found in non-European ancestries by 15% (Figure 2A, 430 regions were replicated with the univariate test, 495 regions were replicated when considering both tests). A substantial number (367) of loci were shared by the European, Asian and trans-ancestry analyses and 10% of those shared associations are detected by the joint test. Multi-trait GWAS can increase the number of replicated associations with no additional samples.

### Similarity of the multi-trait signal between ancestry pairs

To quantify whether ancestries had similar multi-trait association patterns in overlapping significant loci, we compared multi-trait vectors of genetic effects in shared loci across all pairs of ancestry, measured as the squared correlation (R²) of genetic effects across traits (Figures 2B-C). Multi-trait effect size vectors were strikingly similar between the EUR and EAS superpopulations with 75% of SNP pairs in shared loci having a correlation across traits higher than 0.8 (Figure 2C, Figure S14). At first glance, the similarity between the EUR and AFR superpopulations seemed lessened. However most of the low correlation SNPs pairs (80% of the shared loci with a rho <0.8) were located on chromosome 1 near the ACKR1 (Figure 3) locus (Chromosome 1: 159,204,875-159,206,500 forward strand), which is known to be adaptive to overcome the p.vivax induced malaria[8,33]. After accounting for the ACKR1 locus the median of correlation between EUR and AFR significant loci was 0.87. Overall, except for the ACKR1 locus, multi-trait patterns were shared between ancestries for hematological traits.

**Figure 3.**
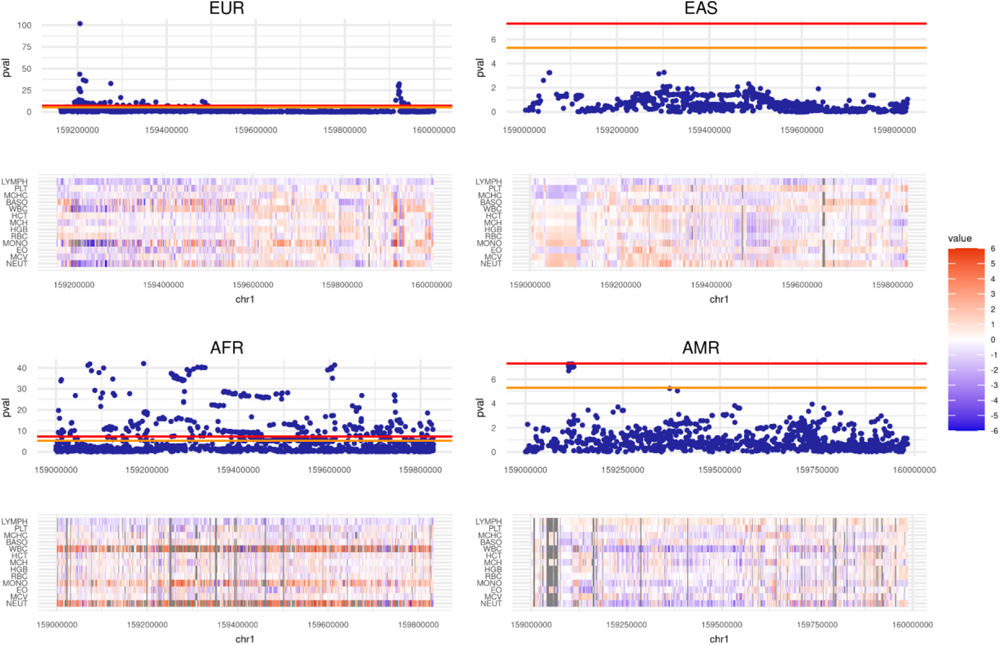
Examples of a diverging multi-trait signal: the ACKR1 locus. For each ancestry, Manhattan plot and corresponding multi-trait signal heatmap for hematological traits at the ACKR1 locus. Under each Manhattan plot, the normalized SNP genetic effects (z-scores) are reported through a heatmap. Colors represent the value of the Z-scores. hematological traits order: LYMPH, NEUT, MCV, EO, MONO, RBC, HGB, MCH, HCT, WBC, BASO, MCHC, PLT, MPV, RDW.

### Functional annotation for hematological trait GWAS

To validate the biological relevance of associations found by the joint test, and in particular novel associations, we focused our functional analysis on hematological traits. Indeed, for hematological traits, the joint test detected enough associations in all ancestries to allow for a comparison of the relevance of the functional enrichment across ancestries. Briefly, we mapped lead SNPs using a combination of positional and eQTLs mapping (**Table S12** reports the number of genes mapped by each method). Lead SNPs were mapped to their nearest genes if their distance was less than 10kb. For eQTLs mapping, we selected eQTLs detected in the immune system and blood cells (**Methods**), which corresponds to the trait assessed in the GWAS. To characterize genes found by the annotation step, we looked in which tissue the genes were differentially expressed (**Figure S17**). For each population, we represented the five first tissues with the highest p-value for upregulated genes (Figure 4A). Blood was significantly enriched for the four populations.

**Figure 4.**
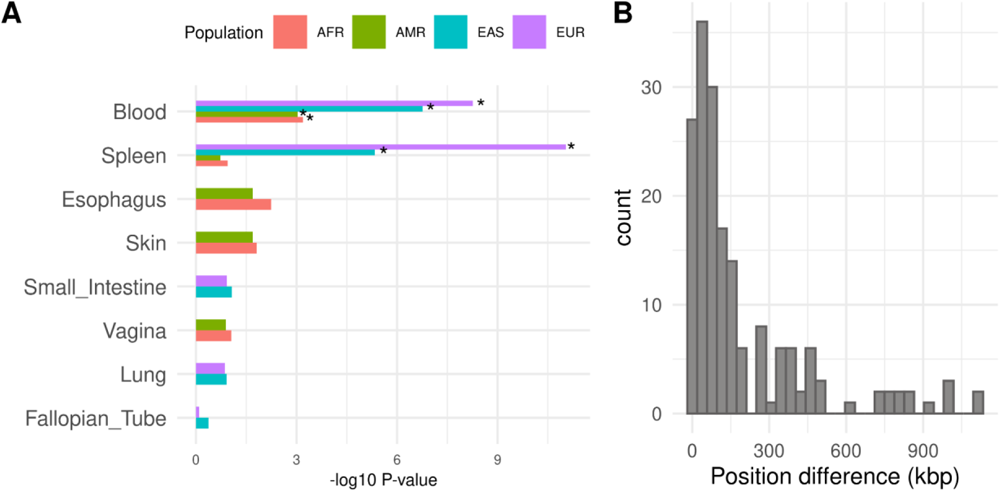
Shared functional features across ancestries. A) Top five tissues with the highest p-value for upregulated genes for the four populations. Stars show significant enrichments (Pbon < 0.05). B) Histogram of the difference between the top SNPs positions in European and East Asian populations. SNP difference positions between European and East Asian populations for 177 genes shared by these two populations.

To further analyze the annotation of the associations discovered by the joint test, we selected genes linked to a SNP significant for the joint test. We then selected genes annotated in GO (Gene Ontology) term sets [34] (**Methods, Tables S13-S16**). For European ancestry, 3 342 genes were identified, 388 genes for the EAS population and 49 genes for the AFR population. For the Admixed American population, there were no genes related to the genetic associations found by the joint test only.

We then investigated whether newly detected signals corresponded to the same regulatory regions across ancestries by computing the position shift between European and East Asian populations for SNPs regulating the same gene. (i.e., SNPs that are eQTLs for the same gene, **Methods, Table S17**). Here, we focused on the European and East Asian ancestries because the two ancestries share enough significant loci (433, Figure 1A) to assess whether these loci systematically regulate the same genes. A total of 177 genes were shared in both ancestries (i.e., associated with a lead SNP through a significant eQTL). When there were several SNPs in one ancestry for one gene, we kept those with the closest pair between populations. The median distance was 86 kb and for 90% of the SNPs, the distance was under 481 kbp (see Figure 4B). This suggests that the same regulatory regions are involved in gene regulation in both populations.

Because of the relatively high proportion of newly associated variants specific to the African ancestry population, we conducted a targeted functional analysis to validate genes associated with newly identified SNPs in that population. 49 genes were associated with a significant lead SNP in the AFR population. Each of those genes was looked up in The Human Gene Database GeneCards (www.genecards.org, [26]) (**Table S17**). Five genes mapped to hemoglobin subunits (*HBG1, HBG2, HBE1, HB*B and *HBD*) involved in beta thalassemia and fetal hemoglobin quantitative trait locus 1 diseases. Eight genes mapped to other blood-related pathways like Tubulin Beta 1 Class VI (*TUBB1*) expressed in platelets and CD36 molecule (*CD36*), a gene protein located at the platelet surface. Five genes mapped to immune system-related pathways. 20 genes mapped to the olfactory receptor family. This overrepresentation of Olfactory receptors reflects probably more the close proximity of these genes along the genome rather than a genuine functional enrichment. Indeed, 15 SNPs mapping to an olfactory receptor family gene are located in a 1 Mbp region (chr11:4.449.477 bp to chr11:5.539.485 bp). For African ancestry, 37% of associations that were discovered by the joint test and annotated with GO term genes mapped to relevant blood pathways.

## Discussion

In this study, we applied JASS, a multi-trait analysis pipeline previously developed and tested on European ancestry to four non-European ancestries. On three (AMR, EAS and AFR) out of four ancestries, our pipeline generalized without barriers and allowed for the detection of new associations while controlling for the genomic inflation factor. We also designed a Transancestry extension of the omnibus test. Altogether, single population and trans-ancestry, omnibus tests identified 367 new associations in non-European ancestry. For hematological traits, these new associations were linked with genes upregulated in blood, a relevant tissue, in all ancestries. Notably for AFR ancestry, a fine analysis of new associations pointed toward blood and immune pathways in 37% of annotated genes. Overall, this study highlights that imputation followed by multi-trait testing can be a valuable tool set for non-European ancestry analysis.

Our extension of the omnibus test to perform trans-ancestry GWAS is an elegant and computationally efficient way to perform trans-ancestry GWAS without assuming homogeneous effects across ancestries. Indeed, the trans-ancestry omnibus test relies on the independence of the chi-squares derived in each ancestry to aggregate them in an unique chisquare test. Hence, in each ancestry, the genetic signals can deviate from the null hypothesis in different directions and contribute to the test statistic. However, our trans-ancestry analysis detected only a handful of new signals, suggesting that the additional degree of freedom in the expected distribution under the null hypothesis dilutes the signal and leads to a lack of statistical power. Incorporating LD local structure of each ancestry into the multi-trait test, as has been done for univariate GWAS[35], might enable us to further improve our trans-ancestry multi-trait tests.

Further improvements of JASS could include a better adaptation to admixed populations. Here, we validated the use of the JASS pipeline for admixed ancestries by showing that the omnibus test is not inflated for Admixed Americans (Figures S10 and S11), and by assessing the accuracy of our estimate of the distribution of Z-scores under the null in the Caribbean population (Figure S4), which is admixed between the African and European ancestries[4]. Although our current approach appears valid and was able to detect 15 new associations in the Admixed Americans, it may not be optimal in terms of statistical power or may be biased for higher degrees of admixture. We recommend a careful inspection of the genomic inflation factor when using JASS on admixed populations. Further development of the method may allow JASS to leverage local ancestry in admixed populations to boost GWAS statistical power as previously done on individual level data [36].

We encountered an unforeseen pitfall with the SAS ancestry: the imputation led to an implausible increase in the number of associations for the joint test. After an investigation, this inflation was due to collinearity of traits investigated coupled to a high sample overlap. This specific set of conditions leads to an increased condition number for covariance under the null, or more plainly said, to a null hypothesis that lacks robustness (Supplementary Note 3). Hence, we do not recommend using the omnibus test in this specific setting. We will further investigate the robustness of the omnibus test on SAS data to confirm that this inflation is not related to other specificities of this population.

A large fraction of new genetic associations matched significant eQTL: 80% (1729/2173) for the European, 60% (163/271) for the East Asian and 82% (59/72) for the African population. However, for the AMR population, functional annotation was less informative suggesting a lower multi-trait or eQTLs annotation signal quality (only 1.9% of sampled individuals in GTEx self-reported as hispanic [37]). Only six significant SNPs out of 57 were eQTLs for a gene. This observation underlines the importance of representing a diverse population not only in GWAS sampling but also in other genomic assays.

Our assessment of multi-trait genetic signal similarity across ancestries demonstrates that the omnibus test allowed for an increased replication of loci detected in European ancestry in non-European ancestries. We mapped associated SNPs to genes through eQTLs and observed that genes share the same regulatory regions in the EUR and EAS ancestries. We reported a systematic comparison of multi-trait genetic signals in shared loci. We observe strikingly similar multi-trait patterns (with the median of the genetic signal squared correlation at variant pairs being above 0.8) except in ACKR1 locus which is tied to an adaptation specific to African ancestries protective for p.vivax parasites.

Our observations suggest an overall stability of the sign of genetic effects except in the ACKR1 loci and are consistent with previous reports that genetic effects are consistent across ancestries [30,38,39]. Other reports nuance the extent of this similarity by noting that genetic correlations across ancestries are often significantly lower than 1 [12,40–42], and loci replication can be lower for specific traits with culturally dependent definition, such as depression (only 11% of European lead SNPs for depression are replicated East Asian ancestry [41]). Hematological traits – continuous traits with an objective definition – can be a good setting to assess replication with less confounding.

Our observation at the ACKR1 loci suggests that modification of multi-trait genetic effects is associated with evolutionary forces. We hypothesize that divergences in multi-trait effects are more striking and more telling functionally than a difference in mean between univariate effect sizes. A systematic investigation coupling trans-ancestry multi-trait genetics and evolutionary pressure measures could highlight how recent evolutionary events in the human population transformed genetic effects. In short, through our multi-trait and trans ancestry GWAS we detected relevant new associations and highlighted the similarity of multi-trait genetics across ancestry. We argue that a computationally efficient pipeline such as the JASS pipeline could be a tool of choice to investigate a multi-trait genetic pattern across ancestries and their potential coupling with evolutionary forces.

## Conclusion

By conducting multi-trait GWAS on 93 summary statistics originating from five ancestries, we detected 367 new genome-wide significant associations in non-European populations (15 in AMR, 72 in AFR and 280 in EAS), which represents respectively 7%, 25% and 21% of all associations in the AFR, AMR and EAS populations. Overall, multi-trait testing increases the replication of European associated loci in non-European ancestry by 15%. Pleiotropic effects were highly similar at significant loci across ancestries (e.g. the mean correlation between multi-trait genetic effect of EUR and EAS was 0.88). For hematological traits, strong discrepancies in pleiotropic effects are tied to known evolutionary divergences: the ARKC1 loci which is adaptive to overcome the p.vivax induced malaria. Altogether these analyses suggest that multi-trait GWAS methods can be a valuable tool to narrow the genetic knowledge gap between European and non-European populations.To facilitate multi-trait GWAS on non European ancestries, we distribute publicly (***Availability of data and material*** section) the JASS pipeline, and curated entry files (summary statistics, Reference panel) issued from this study.

## List of abbreviations

EUR - European

AMR - Admixed American

AFR - African

EAS - East Asian

SAS - South-East Asian

## Declarations

### Ethics approval and consent to participate

Not applicable.

### Consent for publication

Not applicable.

### Availability of data and material

All computational tools used in this study are publicly available on the gitlab of the Statistical Genetics group of the Institut Pasteur:

– JASS pipeline: https://gitlab.pasteur.fr/statistical-genetics/jass_suite_pipeline
– RAISS: https://gitlab.pasteur.fr/statistical-genetics/raiss
– JASS: https://gitlab.pasteur.fr/statistical-genetics/jass
– JASS preprocessing https://gitlab.pasteur.fr/statistical-genetics/jass_preprocessing

Reference panels used in this study can be downloaded at https://gitlab.pasteur.fr/statistical-genetics/jass_suite_pipeline/-/tree/master/Ref_Panel

LDscores for the five ancestries can be download at: https://doi.org/10.5281/zenodo.8096588

Linkage disequilibrium matrices for the five ancestries can be downloaded at: https://doi.org/10.5281/zenodo.10391372

Summary statistics imputed and harmonized by the JASS pipeline can be downloaded on the Zenodo platform for each ancestry under the following doi:

AFR - https://doi.org/10.5281/zenodo.8060264

EAS - https://doi.org/10.5281/zenodo.8068881

AMR - https://doi.org/10.5281/zenodo.8068935

EUR - https://doi.org/10.5281/zenodo.8068972

TRANS - https://doi.org/10.5281/zenodo.10213745

Script and data to reproduce Figures presented in this manuscript: https://zenodo.org/records/10299388

## Funding

This research was supported by the Agence Nationale pour la Recherche (ANR-20-CE36-0009-02). This work has been conducted as part of the INCEPTION program (Investissement d’Avenir grant ANR-16-CONV-0005)

## Authors’ contributions

HJ conceptualized the study, supervised the analysis. LT and LH performed the data curation, the formal analysis. LT, HJ and DF investigated the results. HJ and LT wrote the original manuscript. HJ, LT, DF and HA reviewed the original draft. LT and HJ prepared visualisation of results. HJ and HA acquired funding for this work. All authors read and approved the final manuscript.

## Acknowledgements

We would like to thank members of the Statistical Genetic Unit at the Institut Pasteur who contributed to the quality of this study through their insightful feedbacks, namely: Antoine Auvergne, Christophe Boetto, Jonas Lefevre-Daumal, Arthur Frouin, and Yuka Suzuki.

This research has been conducted using the UK Biobank Resource under Application Number 42260.

## Competing interest

The authors declare that they have no competing interests.

## Notes

https://gitlab.pasteur.fr/statistical-genetics/jass_suite_pipeline

https://gitlab.pasteur.fr/statistical-genetics/raiss

https://gitlab.pasteur.fr/statistical-genetics/jass

https://gitlab.pasteur.fr/statistical-genetics/jass_preprocessing

https://doi.org/10.5281/zenodo.8060264

https://doi.org/10.5281/zenodo.8068881

https://doi.org/10.5281/zenodo.8068935

https://doi.org/10.5281/zenodo.8068972

https://doi.org/10.5281/zenodo.8096588

https://doi.org/10.5281/zenodo.10391372

### Competing Interest Statement

The authors have declared no competing interest.

### Summary of Updates

● We clarified the motivation for our study in the Introduction and in the Overview of the study sections. ● We conducted an additional validation analysis to ensure that key parameters underlying the JASS association test could be accurately inferred on admixed populations. ● We computed and reported the number of independent loci detected exclusively with the univariate tests. ● We discussed the gain in number of associations provided by the joint test more extensively. ● We provided an archive containing all data and scripts necessary to reproduce Figures presented in the manuscript. ● We edited the functional analysis part of the paper to clarify its scope and aim.

