## Supplementary Notes and Figures for "Multi-trait GWAS for diverse ancestries: Mapping the knowledge gap"

#### Table of contents


Additional file legends 1
Supplementary Note 1: Extending JASS analysis pipeline to non-European ancestries 2
Supplementary Note 2: Hematological trait heritabilities and genetic correlations across populations 2
Supplementary Note 3: Highly correlated Z-scores under the null coupled to imputation lead to an inflation of the multi-trait test 3
Supplementary Figures 5


#### Additional file legends

**Additional file 1: Table S1** - Ancestries used in recently published methodological developments in Statistical genetics

**Additional file 2: Table S2** - GWAS Summary Statistics considered for each superpopulation

**Additional file 3: Table S3** - RAISS performances metrics (across ancestry and traits) for RAISS imputation for the chosen parameters (--R2-threshold 0.6, --minimum-ld 5 and --eigen-threshold 0.05) measured on the chromosome 22

**Additional file 4: Table S4** - Covariances of Zscores under the null hypothesis (no genetic effects)

**Additional file 5: Table S5** - Genetic covariance matrices for the four studied superpopulations

**Additional file 6**: Independent Significant SNPs detected by the Omnibus test. The column 'Discovered by' indicates if the SNPs was identified only by the joint test. SNPs annotated 'Joint only' are identified by multivariate test only and annotated 'Both' are identified by univariate and joint tests. **Table S6** - SNPs associated with hematological traits and glycemic traits in EAS ancestry. **Table S7 -** SNPs associated with hematological traits and glycemic traits in AMR ancestry. **Table S8** - SNPs associated with hematological traits and glycemic traits in AFR ancestry. **Table S9** - SNPs associated with hematological traits and glycemic traits in EUR ancestry. The column 'Discovered by' indicates if the SNPs was identified only by the joint test. **Table S10** - SNPs associated with hematological traits in the Trans-ancestry analysis.

**Additional file 7: Table S11**- New associations (significant for the Joint test only) detected for hematological traits for all ancestries with their allele frequency in each population

**Additional file 8: Table S12** - Number of genes mapped to SNPs associated to hematological traits with the joint Test

**Additional file 9**: Gene Ontology term enrichment across ancestries. **Table S13** - Gene Ontology term enrichment for all genetic association found in EAS population with FUMA GWAS. **Table S14** - Gene Ontology term enrichment for all genetic association found in AMR population with FUMA GWAS. **Table S15** - Gene Ontology term enrichment for all genetic association found in AFR population with FUMA GWAS. **Table S16** - Gene Ontology term enrichment for all genetic association found in EUR population with FUMA GWAS.

**Additional file 10: Table S17** - For each gene, pairs of eQTLs associated in European and East asian ancestry and their position difference

**Additional file 11: Table S18** - Functional annotation of genes associated with SNPs detected by the joint test only in AFR population.

**Additional file 12**: Supplementary notes and Figures

#### Supplementary Note 1: Extending JASS analysis pipeline to non-European ancestries

We previously implemented a nextflow pipeline to harmonize data and to perform multi-trait GWAS using JASS[1]. While the JASS pipeline could be applied on any ancestries provided reference panels and input files in the proper format, we provided precomputed ones only for European ancestry. To facilitate the use of JASS on diverse ancestries, reference panels for the 5 superpopulations defined in 1000G[2] and other required input files can be downloaded through the following link: https://gitlab.pasteur.fr/statistical-genetics/jass_suite_pipeline.

#### Supplementary Note 2: Hematological trait heritabilities and genetic correlations across populations

Hematological traits have an heritability ranging from low for basophil to high for platelet volume (Figure S3.A). While we acknowledge that the precision of heritability estimates varies by ancestry due to the large discrepancies in sample size (see Table S2), we report heritability estimates when they exceed their standard deviation. As previously reported [3], we noticed significant heritability discrepancies between populations. Interestingly, genetic correlation highlights the different interpretation of classical blood assays between ancestries (Figure S3.B). In the European, East Asian and American population, the gold standard for diabetes diagnosis, that is HbA1c, has its highest genetic correlation with fasting glucose (0.32, p-value = 6.67x10-13; 0.66, p-value=0.1; 0.84, p-value= 0.1533). In contrast, in the African population, the most highly correlated trait with HbA1c is the Red blood cell distribution width (RDW=0.88, p-value=0.07). Red blood cell count is modified in individuals with sickle cell anemia and sickle cell traits. While the genetic correlation estimates are not significant except for the European population, this tendency reflects the inadequacy of the HbA1c as a measure of glycemic regulation in parts of the African population.

#### Supplementary Note 3: Highly correlated Z-scores under the null coupled to imputation lead to an inflation of the multi-trait test

For the SAS ancestry specifically, we observed an increase of the number of significant hits for the joint test after imputation (almost twice the number of hits observed before imputation, see table below) for the multi-trait test that was inconsistent with the proportion of imputed SNPs.

| | SAS | AFR | EAS | AMR | EUR |
|---|---|---|---|---|---|
| Number of hit before imputation | 51 | 881 | 1043 | 42 | 14629 |
| Number of lead after imputation | 95 | 947 | 1252 | 57 | 16326 |
| percentage of increase | 86 % | 7.5% | 20% | 35.7% | 11% |
| percentage of imputed SNPs | 17% | 6% | 27% | 22% | 13% |
| Condition number of the H0 matrix | 22.1 | 15.8 | 8.0 | 13.5 | 13.8 |

As the imputation performances and sanity checks, that is imputation did not lead to an increase of significant hits for the univariate test, for this ancestry were comparable to other ancestry, we hypothesize that the inflation was a consequence of a specific interaction between the imputation and the joint test.

To test this hypothesis, we generated Z-scores vector under the null (following a multivariate distribution with mean 0 and covariance matrix given by the intercept of the LDscore regression) and introduce 20% of imputed SNPs by adding a random imputation error coherent with the one observed in our data: normally distributed noise with a standard deviation of 0.1 (comparable to the median imputation error reported in Table S3). We report the distribution of the p-value of the omnibus test for these Z-scores across the 5 ancestry (see Figure S4) and notice a large inflation specific to the SAS ancestry.

We investigated the level of collinearity of the covariance matrices of Z-score under the null ( $\Omega^r$). As derived in [4], for a pair of trait i and j the expected covariance under the null is equal to

$\Omega^r_{ij} = \rho_{ij} \frac{n_s}{n_i n_j}$, where $\rho_{ij}$ is the total covariance between traits $i$ and $j$, $n_s$ the number of samples shared between study $i$ and $j$, and $n_i$ and $n_j$ the sample sizes of study $i$ and $j$. Hence, the covariance is impacted by one factor that is relatively constant across ancestries (the observed covariance between the two traits), and by one other factor that strongly varies depending on the meta-analysis composition: sample overlap between studies. Concerning the SAS ancestry, all samples for hematologic traits are from Ukbiobank (a total sample overlap).

To assess the impact of sample overlap on collinearity, we computed the condition number for the $\Omega^T$ matrix in each ancestry (see table above). The condition number is the square root of the ratio of the largest eigenvalue and the lowest eigenvalue of the covariance matrix. We observe the highest condition number for the SAS ancestry, confirming the higher colinearity of the covariance under the null for this ancestry. In this specific setting – highly correlated phenotypes coupled to a total sample overlap – the omnibus test lacks robustness. Hence we decided to exclude the SAS ancestry from subsequent analysis.

### Supplementary Figures

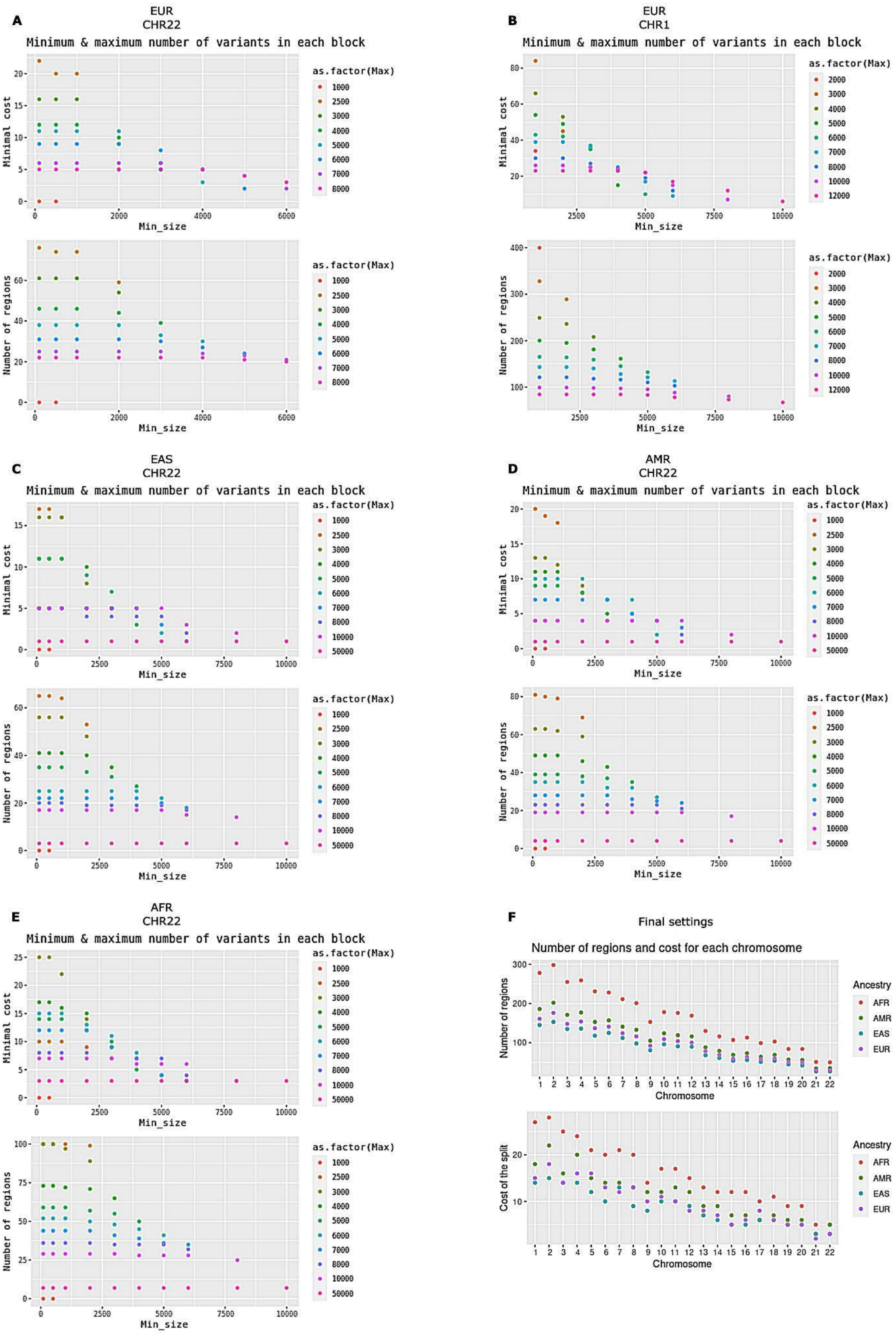


**Figure S1. Cost of the LD-independent blocks proposed by the snp_ldsplit function of the BigSNPr package as a function of the min_size and max_size parameters**. The cost represents the sum of squared correlations outside the blocks. (A-E) variation of the cost (top panel) and of the number of regions in the partition (bottom panel) as a function of the allowed minimum number of SNPs in the blocks (x-axis) and of the maximum number of SNPs in the block (dot colors). (F) Number of Regions and the cost for each chromosome in the four super populations with the chosen settings (thr_r2 = 0.05, min_size = 4000, max_size = 5000, max_K = 400).

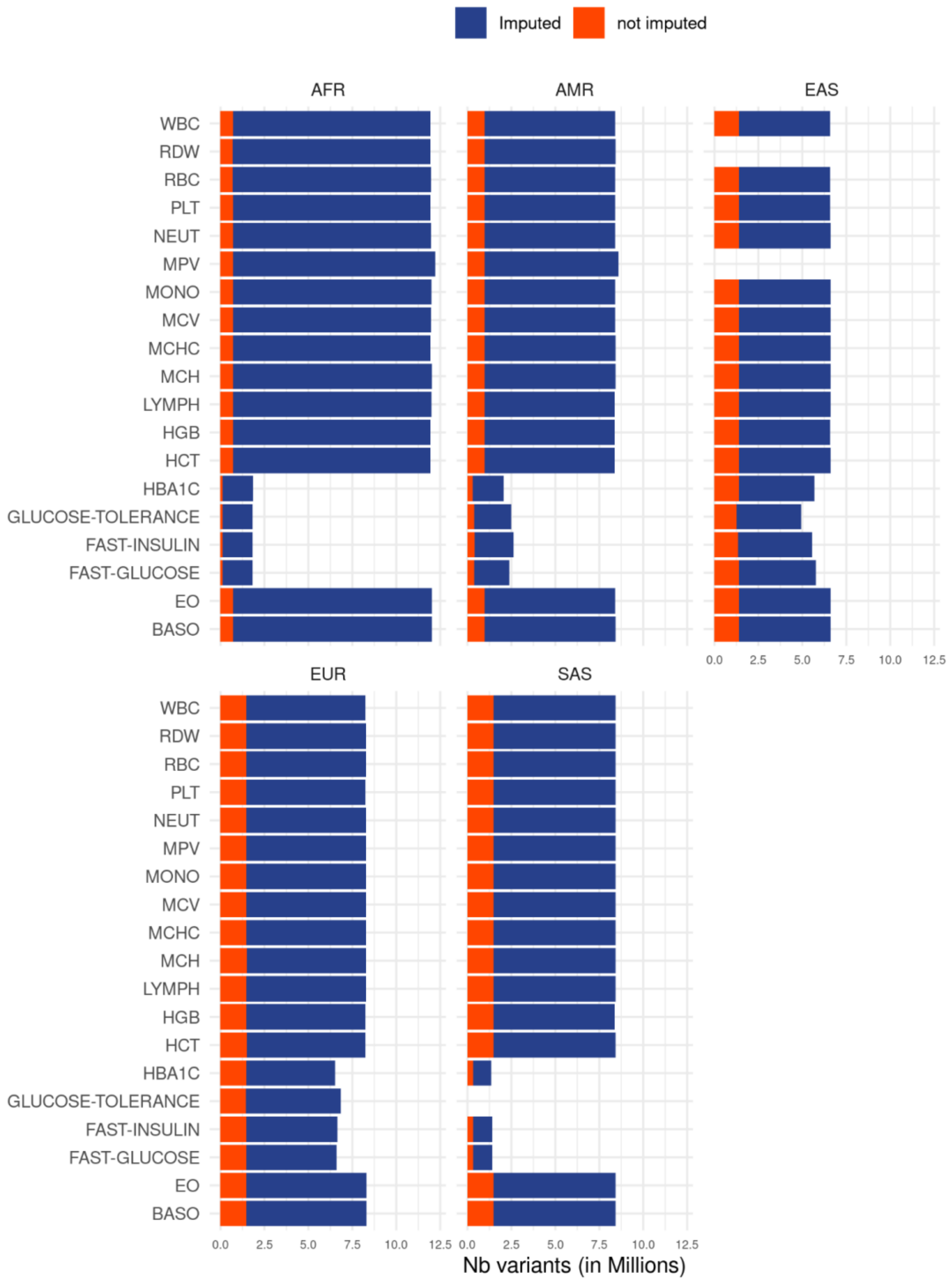


**Figure S2. Barplot of  the number of SNPs by ancestry and traits.** The color represents whether the SNPs have been imputed or not. Orange are Imputed SNPs and Dark blue Snps were present in original data and the reference panel.

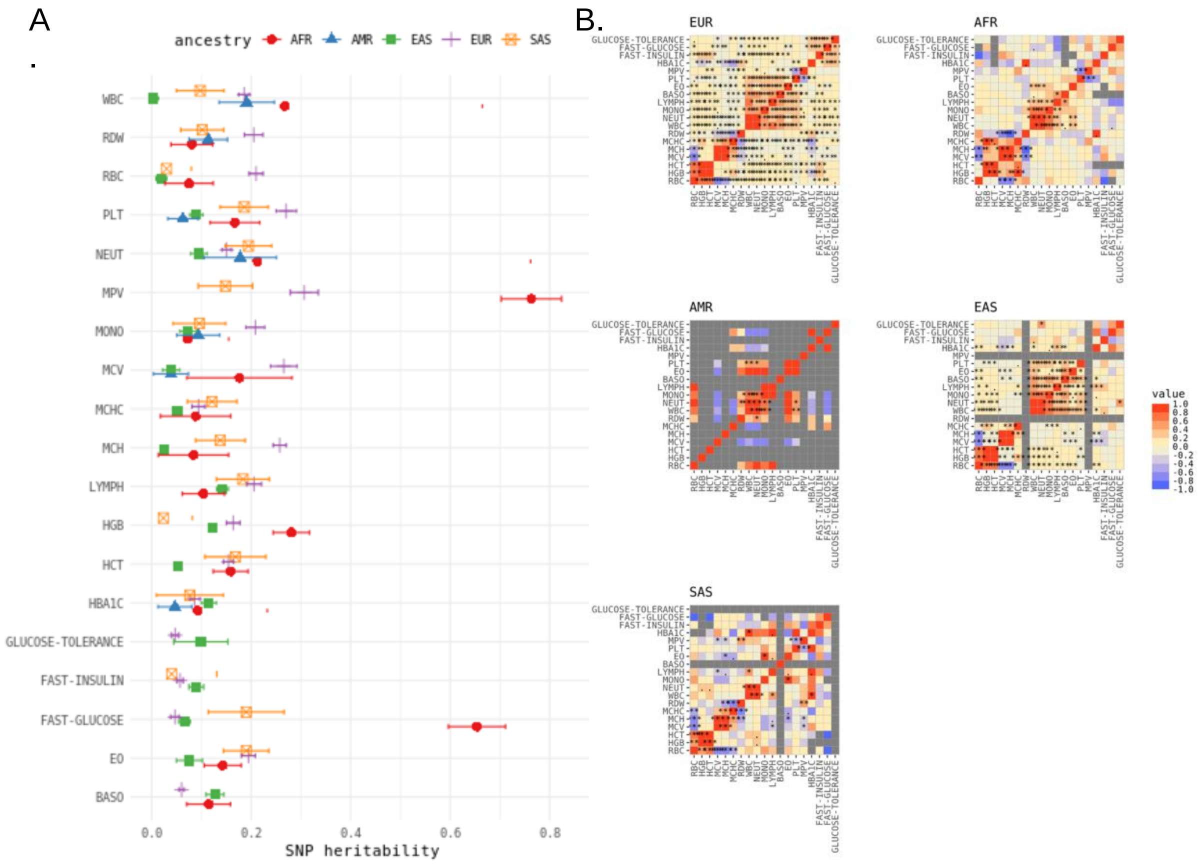


**Figure S3. Heritability estimates and genetic correlation estimates by ancestry.** A) Heritability estimates, colors indicate ancestry and error bars are the standard deviation. B) Genetic correlation between pairs of traits by ancestry on 19 traits.

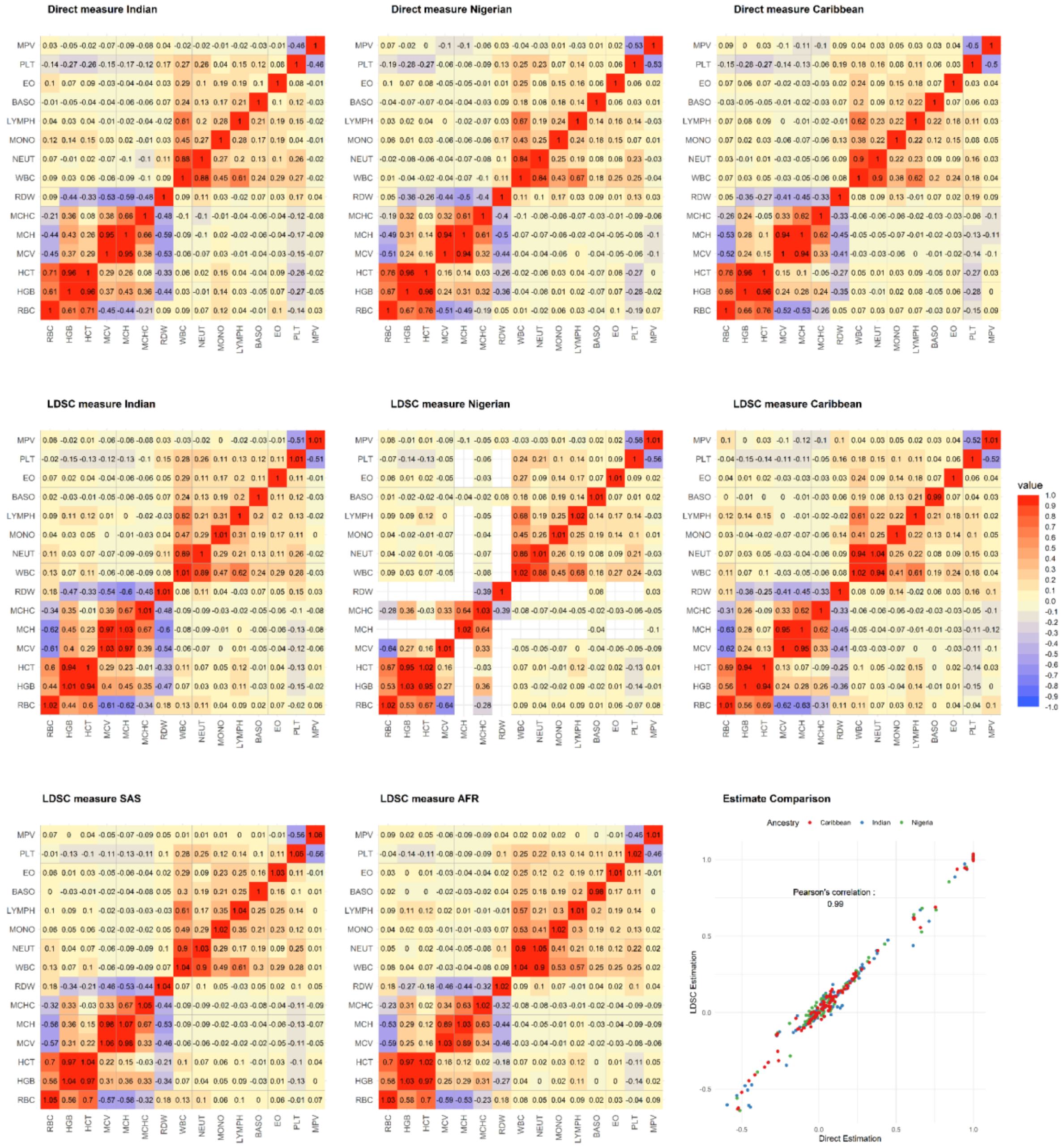

**Figure S4. Comparison of direct and LD-score regression estimations of the Pearson correlations between hematological traits.** Matrices on the first line depict direct estimates of the Pearson correlation between hematological traits in UKBiobank for individuals from the Indian, Nigerian and Caribbean ancestries. The second line displays the same correlation estimated from summary statistics derived in the same individuals. LD-score regression was applied on these summary statistics to derive correlation estimates. The third line displays the covariance under the null for the SAS and AFR ancestry used in this study, and a scatter plot of LDscore correlation estimates with respect to direct estimate of the correlation.

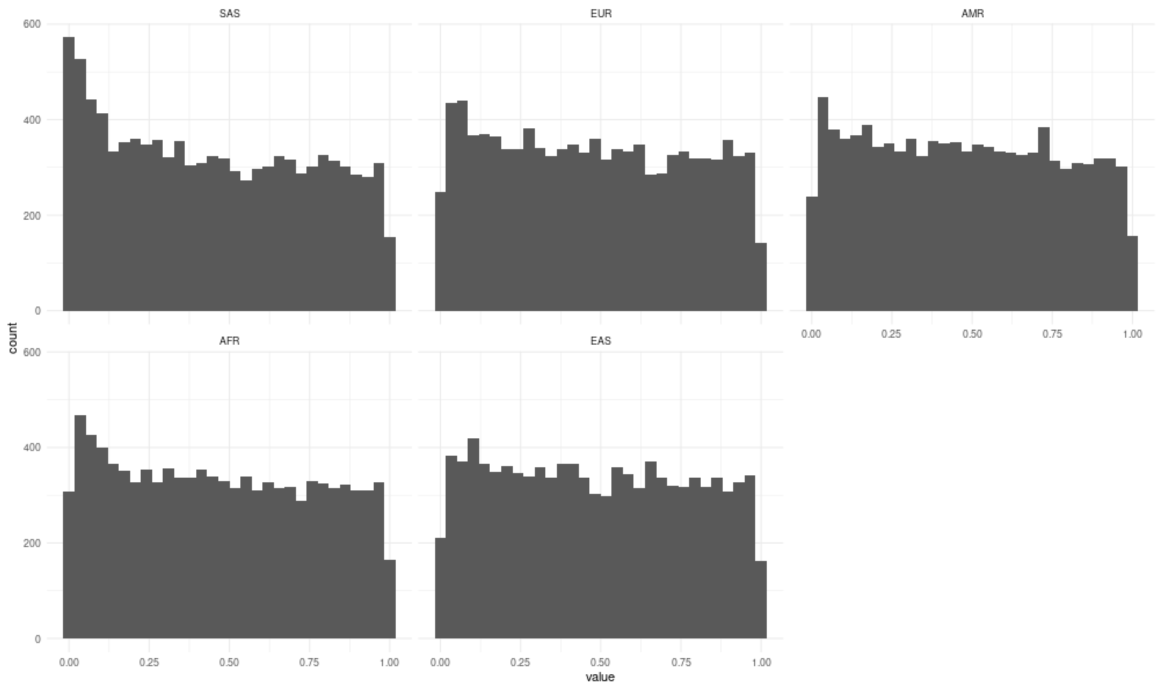


**Figure S5** Distribution of p-value under the null hypothesis for the five ancestries initially considered (SAS, EUR, AMR, AFR, EAS) with data simulated to mimic imputation error.

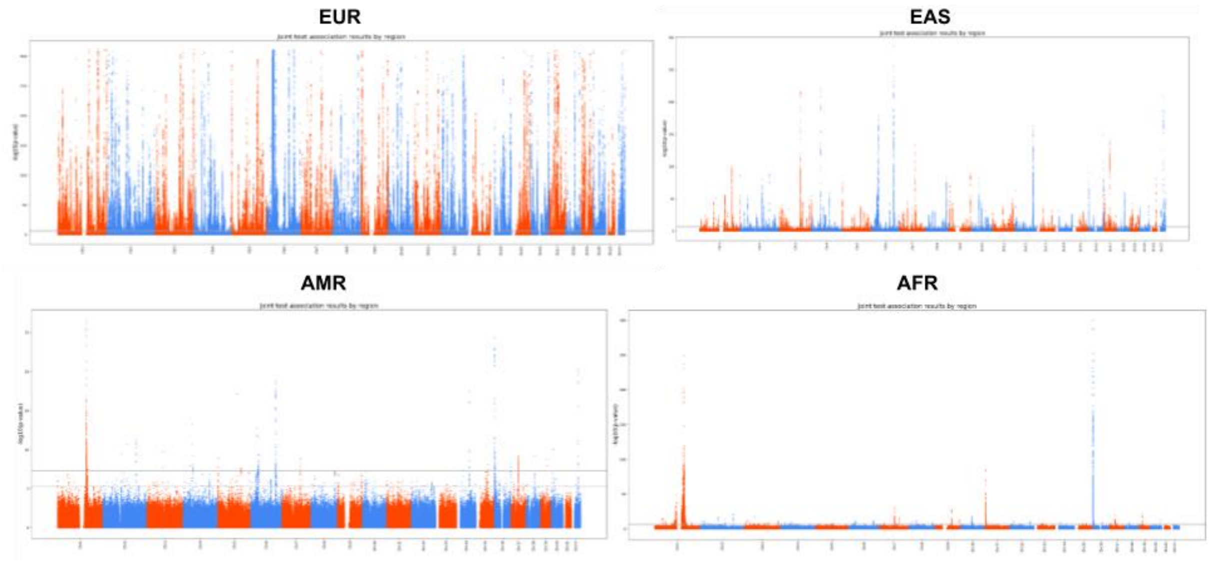


**Figure S6. Manhattan Plots of Multi-trait GWAS for hematological traits in each population.** The dashed line represents 5 x $10^{-6}$ threshold and the solid line represents 5 x $10^{-8}$ threshold.

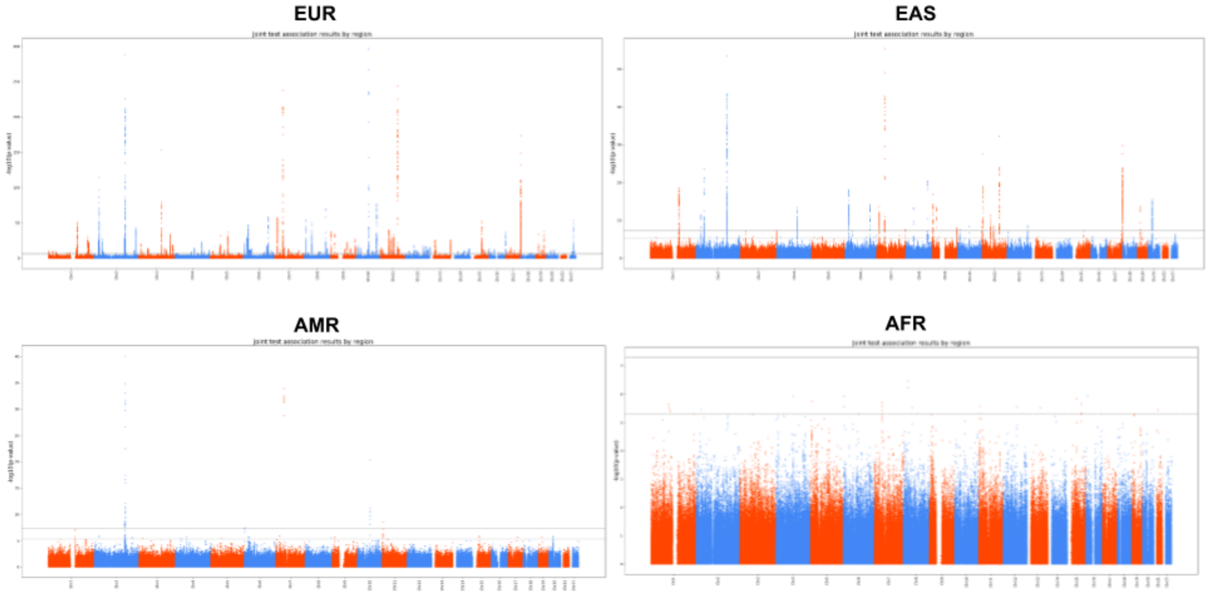


**Figure S7. Manhattan Plots of Multi-trait GWAS for glycemic traits in each population.** The dashed line represents 5 x 10⁻ ⁶ threshold and the solid line represents 5 x 10⁻ ⁸ threshold

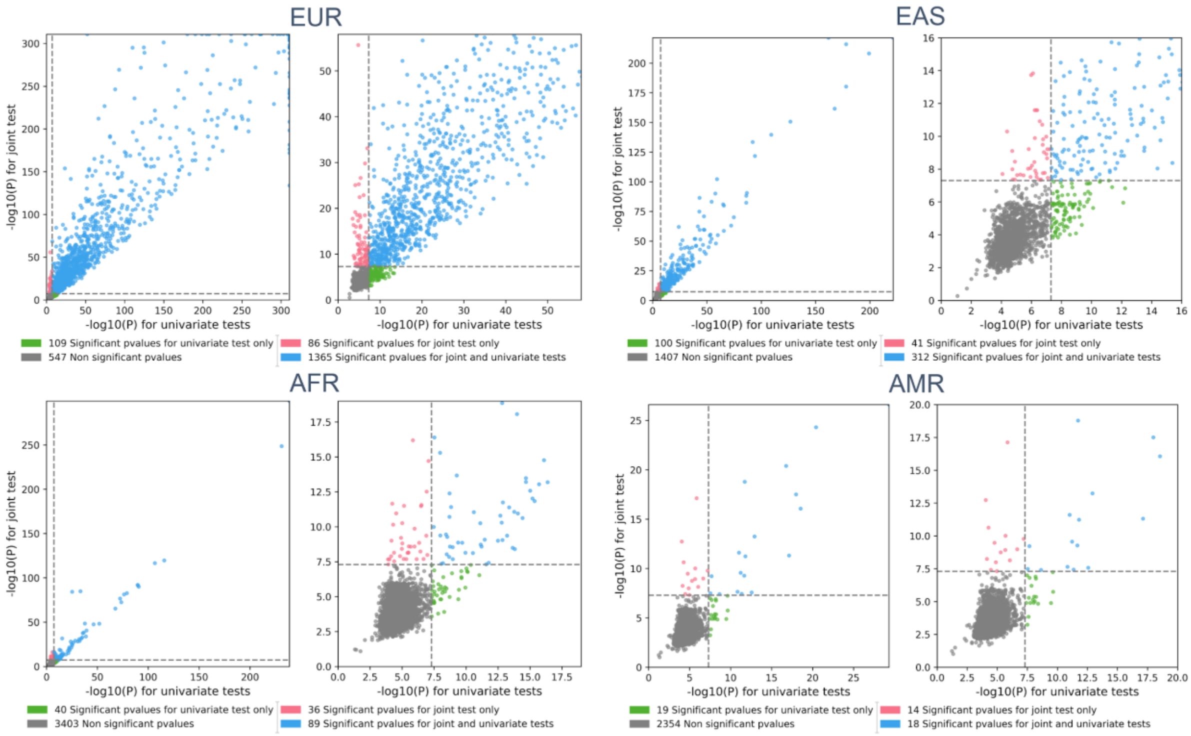


**Figure S8. Hematological trait GWAS Quadrant plots for each ancestry.** The y-axis represents the -log10(*p*-value) for the most significant SNP per region for the *omnibus* test with respect to -log10(*p*-value) for the most significant SNP per region across all univariate GWAS. Complete results are presented in the left sub-panel, and a zoom around the genome-wide significance threshold is presented on the right sub-panel.

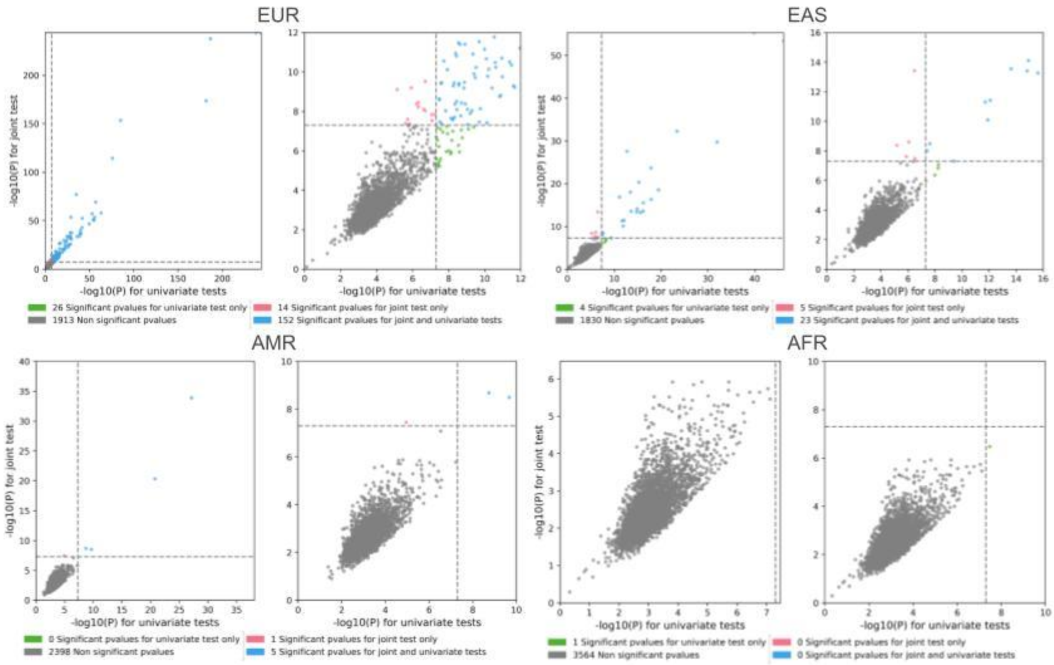


**Figure S9. Glycemic trait GWAS Quadrant plots for each ancestry.** The y-axis represents the -log10(*p*-value) for the most significant SNP per region for the *omnibus* test with respect to -log10(*p*-value) for the most significant SNP per region across all univariate GWAS. Complete results are presented in the left sub-panel, and a zoom around the genome-wide significance threshold is presented on the right sub-panel.

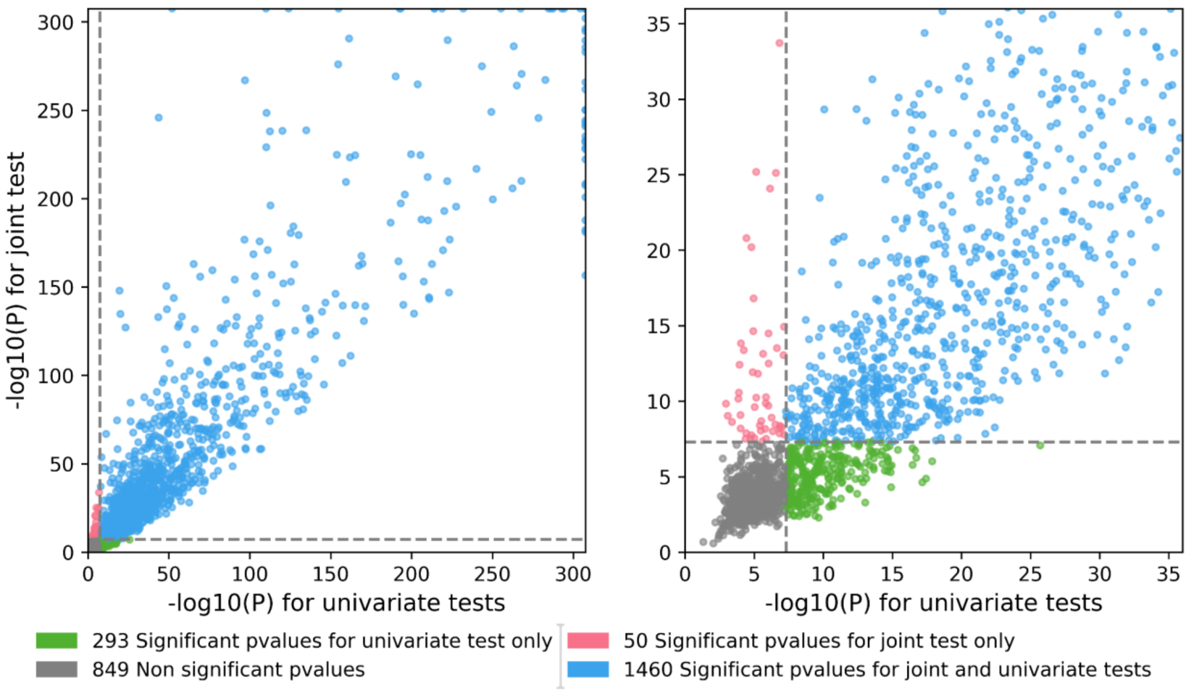


**Figure S10. hematological trait GWAS Quadrant plots for TRANS ancestry.** The y-axis represents the -log10(*p*-value) for the most significant SNP per region for the *omnibus* test with respect to -log10(*p*-value) for the most significant SNP per region across all univariate GWAS. Complete results are presented in the left sub-panel, and a zoom around the genome-wide significance threshold is presented on the right sub-panel.

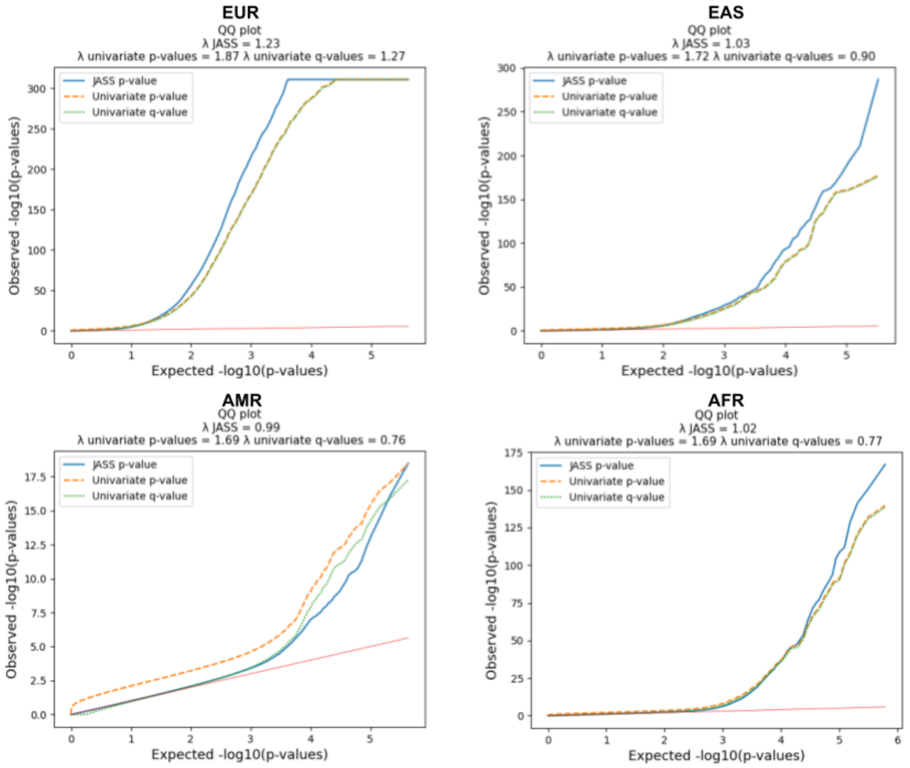


**Figure S11. QQ-PLOT of the p-value of the joint analysis of hematological traits for each population.**

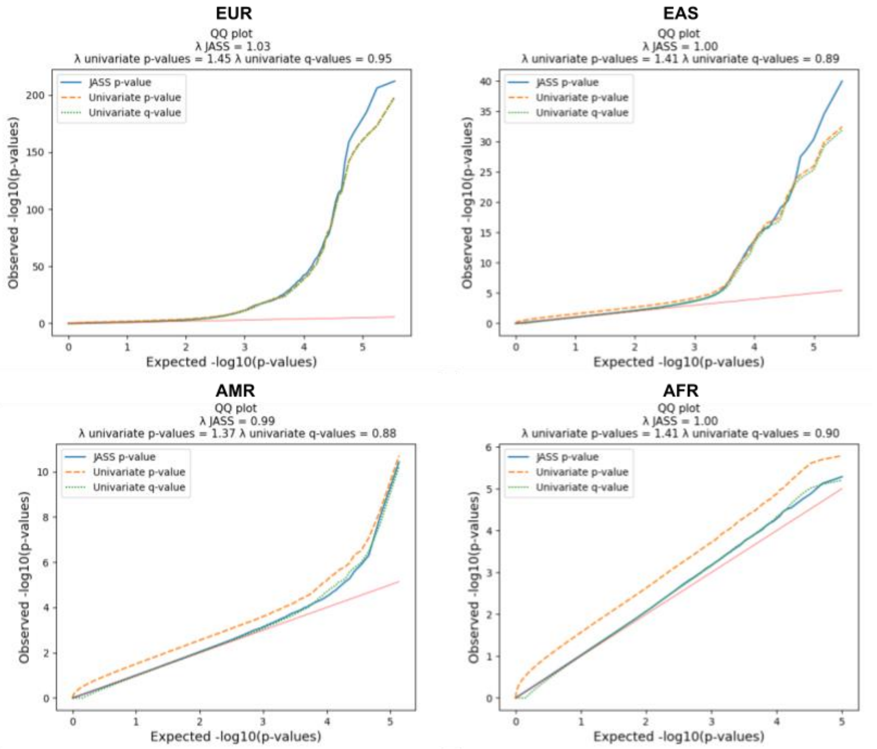


**Figure S12. QQ-PLOT of the p-value of the joint analysis of glycemic traits for each population.**

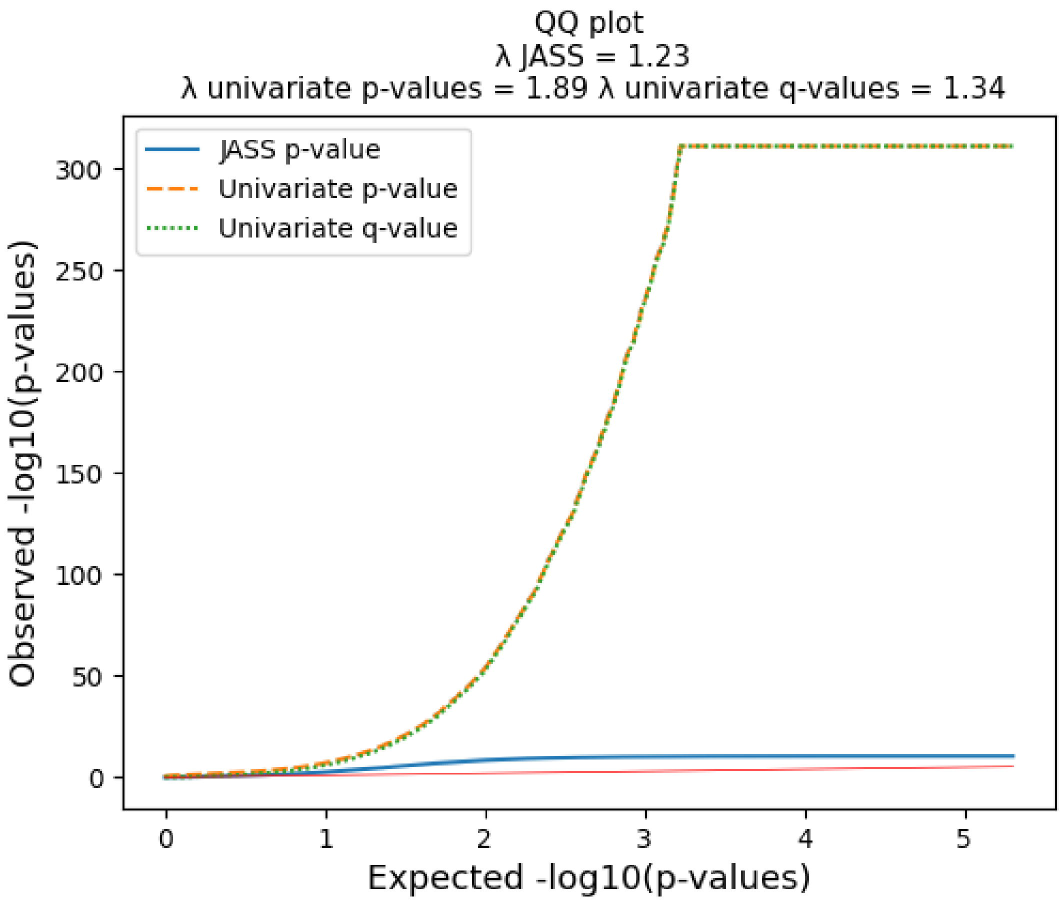


**Figure S13. A QQ-PLOT of the p-value of the joint analysis of blood traits for the TRANS ancestry analysis**

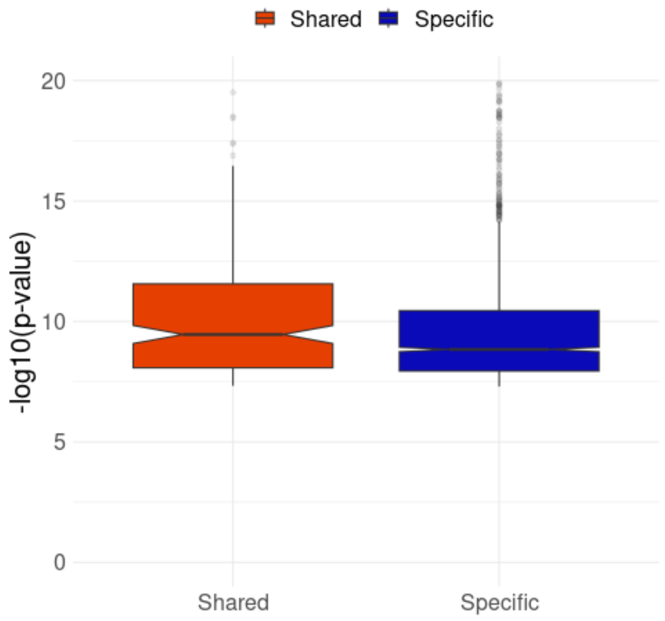

**Figure S14. Distribution of the omnibus test p-value for European significant loci depending on whether they are shared with another ancestry or specific to European Ancestry.**

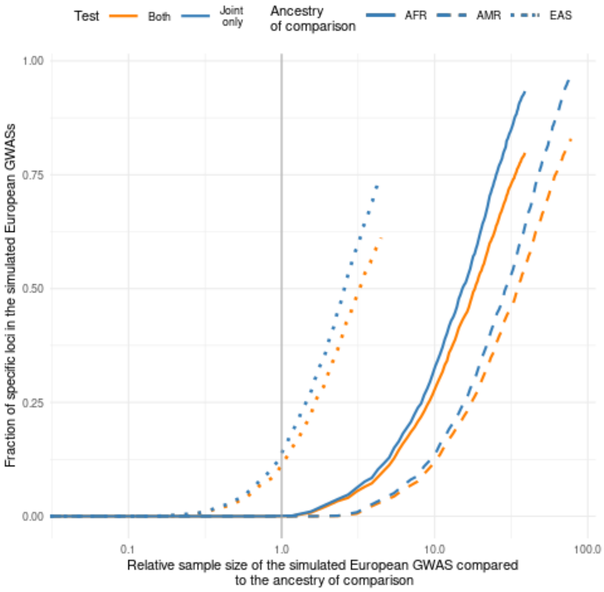


**Figure S15 Fraction of significant European loci that remain specific (i.e. found only in Europe) after simulating a diminished sample size for the European GWASs**. We compared the simulated European multi-trait GWAS on hematological traits to the corresponding analysis in the AMR, EAS and AFR ancestry at their full sample size. Each linetype corresponds to one non-European ancestry. The y axis represents the fraction of loci detected in the simulated European GWASs that were absent in the ancestry of comparison. The x axis is the relative sample size of the simulated European GWAS with respect to the GWAS of comparison.

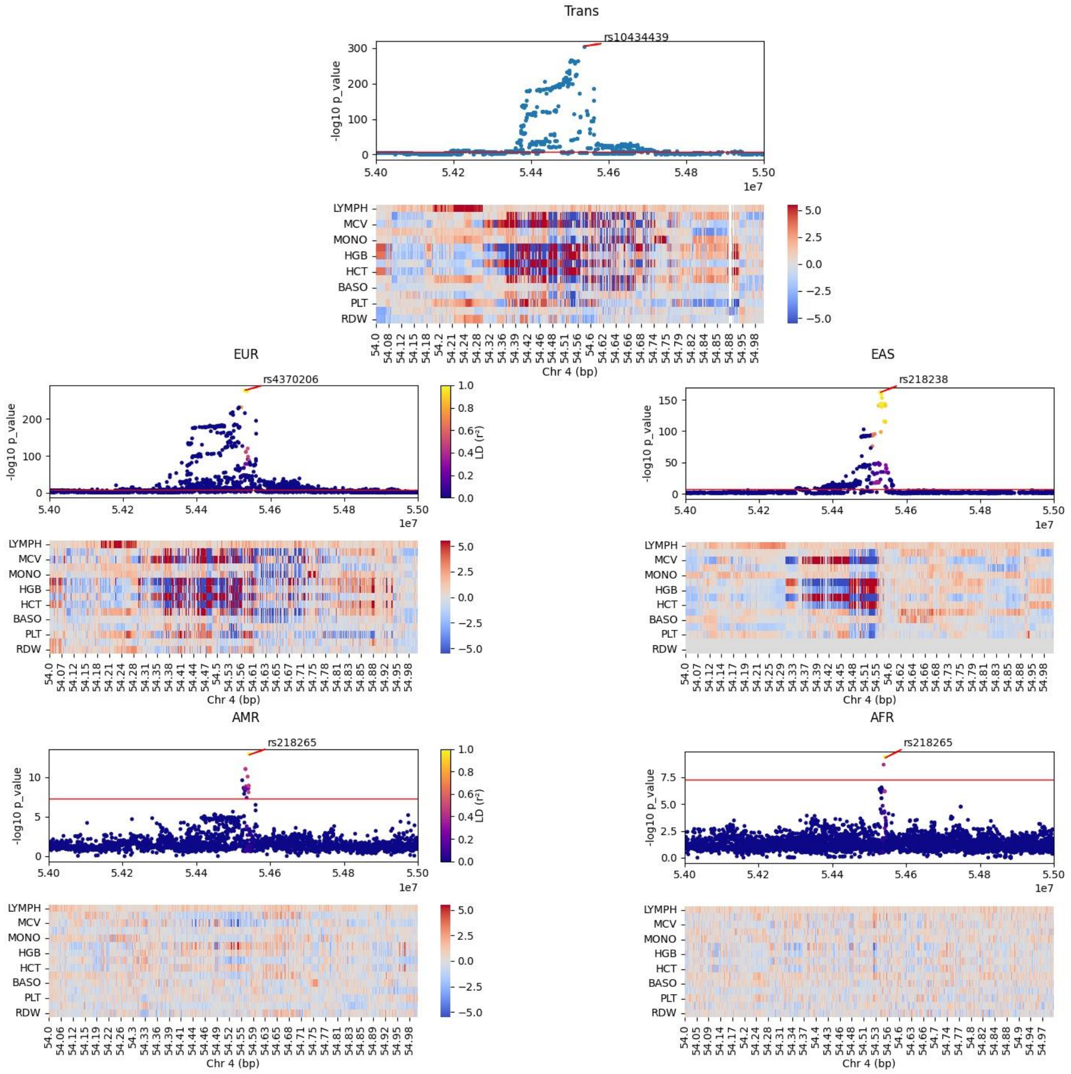


**Figure S16. Manhattan plot and corresponding multi-trait signal heatmap for the TRANS and all ancestry multi-trait GWASs with p-value from univariate test. For ancestry specific GWAS, dots' color represent the LD with the smallest p-value lead SNPs in each panel.** Under each Manhattan plot, the normalized SNPs genetic effects (z-score) are reported through a heatmap. hematological traits order: LYMPH, NEUT, MCV, EO, MONO, RBC, HGB, MCH, HCT, WBC, BASO, MCHC, PLT, MPV, RDW.

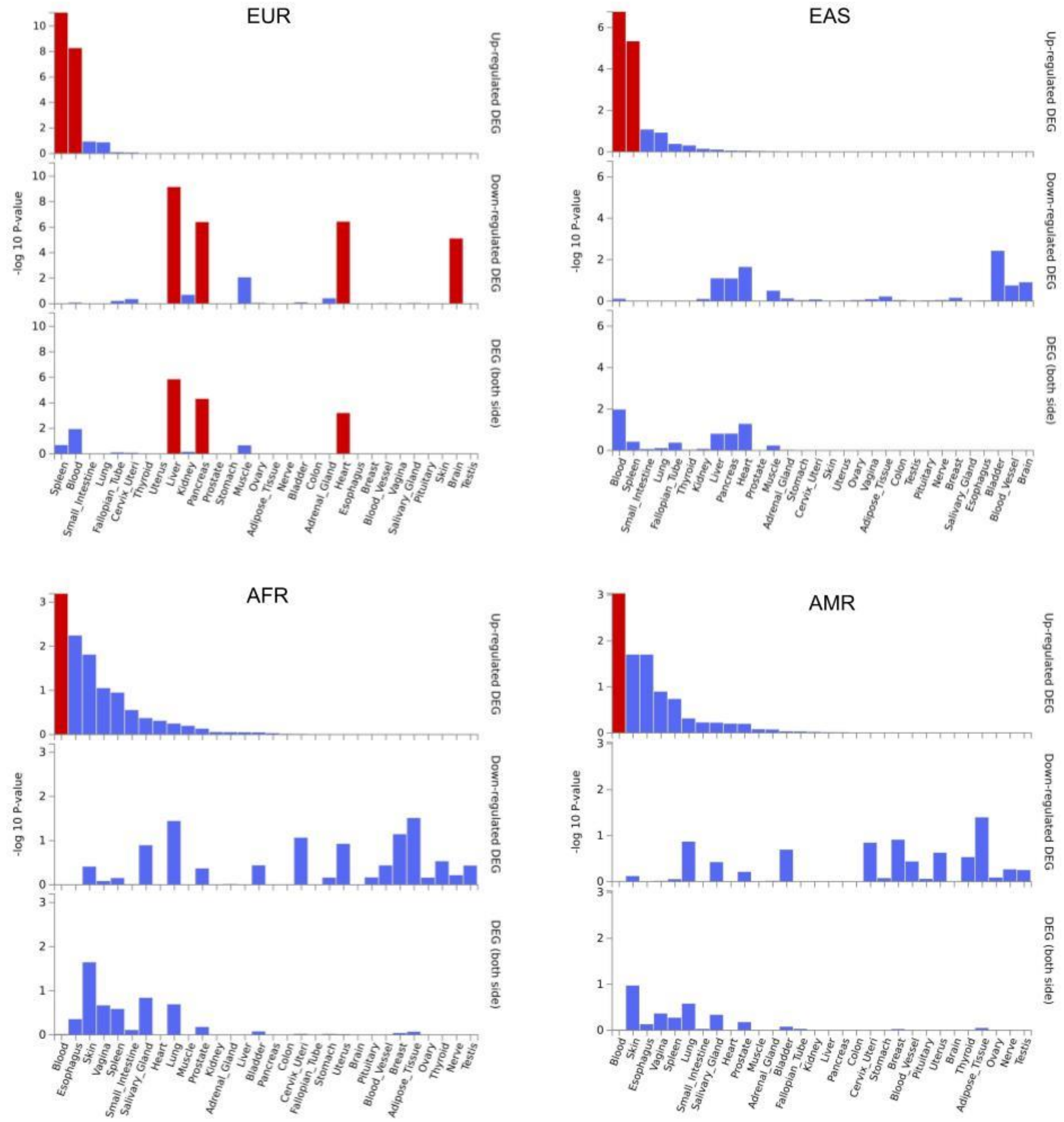


**Figure S17. Differentially expressed genes in four populations**. Differentially expressed genes using GTEx v8 30 general tissue types. Significantly enriched DEG sets ($P_{bon} < 0.05$) are highlighted in red. Tissues are ordered by up-regulated DEG P-value.
